# Unfolding the Apoptotic Mechanism of Antioxidant Enriched-Leaves of *Tabebuia pallida* in EAC Cells

**DOI:** 10.1101/2021.01.11.426226

**Authors:** Md. Mahbubur Rahman, Muhammad Ali Khan, A. S. M. Ali Reza, Khaled Mahmud Sujon, Rokshana Sharmin, Mamunur Rashid, Md. Golam Sadik, Md. Abu Reza, Toshifumi Tsukahara, Ashik Mosaddik, Glenda C Gobe, AHM Khurshid Alam

**Author notes:** **Correspondence:** (1) Prof AHM Kurshid Alam; (2) Professor Glenda C Gobe. **Email addresses of authors:** (M.M.R.); (M.A.K.); (A.S.M.A.R.); (G.C.G); (R.S.); (M.R.); (M.G.S.); (M.A.R.); (K.M.S); (T.T.); (A.M.); (A.H.M.K.A.).

## Abstract

Targeting apoptosis is a promising approach to inhibit the abnormal cell proliferation of cancer progression. Existing anti-apoptotic drugs, many derived from chemical substances, have often failed to combat cancer development and progression. Therefore, identification of apoptosis-inducing anticancer agents from plant-derived sources has become a key aim in cancer research. The present study was designed to explore the regulation of apoptosis by *Tabebuia pallida* (*T. pallida*) using an Ehrlich Ascites Carcinoma (EAC) mouse model and compositional analysis by LC-ESI-MS/MS. Dried and powdered *T. pallida* leaves (TPL), stem bark (TPSB), root bark (TPRB) and flowers (TPF) were extracted with 80% methanol. Using cultured EAC cells and EAC-bearing mice with and without these extracts, anticancer activities were studied by assessing cytotoxicity and tumor cell growth inhibition, changes in life span of mice, and hematological and biochemical parameters. Apoptosis was analyzed by microscopy and expression of selected apoptosis-related genes (Bcl-2, Bcl-xL, NFκ-B, PARP-1, p53, Bax, caspase-3 and -8) using RT-PCR. LC-ESI-MS analysis was performed to identify the major compounds from the most active extracts. In EAC mice compared with untreated controls, the TPL extract exhibited the highest cytotoxicity with significant tumor cell growth inhibition (p< 0.001), reduced ascites by body weight (p< 0.01), increased the life span (p<0.001), normalized blood parameters (RBC/WBC counts), and increased the levels of superoxide dismutase and catalase. TPL-treated EAC cells showed apoptotic characteristics of membrane blebbing, chromatin condensation and nuclear fragmentation, and caspase-3 activation, compared with untreated EAC cells. Moreover, annexin V-FITC and propidium iodide signals were greatly enhanced in response to TPL treatment, indicating apoptosis induction. Pro- and anti-apoptotic signaling after TPL treatment demonstrated up-regulated p53, Bax and PARP-1, and down-regulated NFκ-B, Bcl-2 and Bcl-xL expression, suggesting that TPL shifts the balance of pro- and anti-apoptotic genes towards cell death. LC-ESI-MS data of TPL showed a mixture of glycosides, lapachol, and quercetin antioxidant and its derivatives that were significantly linked to cancer cell targets. In conclusion, the TPL extract of *T. pallida* possesses significant anticancer activity. The tumor suppressive mechanism is due to apoptosis induced by activation of antioxidant enzymes and caspases and mediated by a change in the balance of pro- and anti-apoptotic genes that promotes cell death.

**Graphical Abstract:** 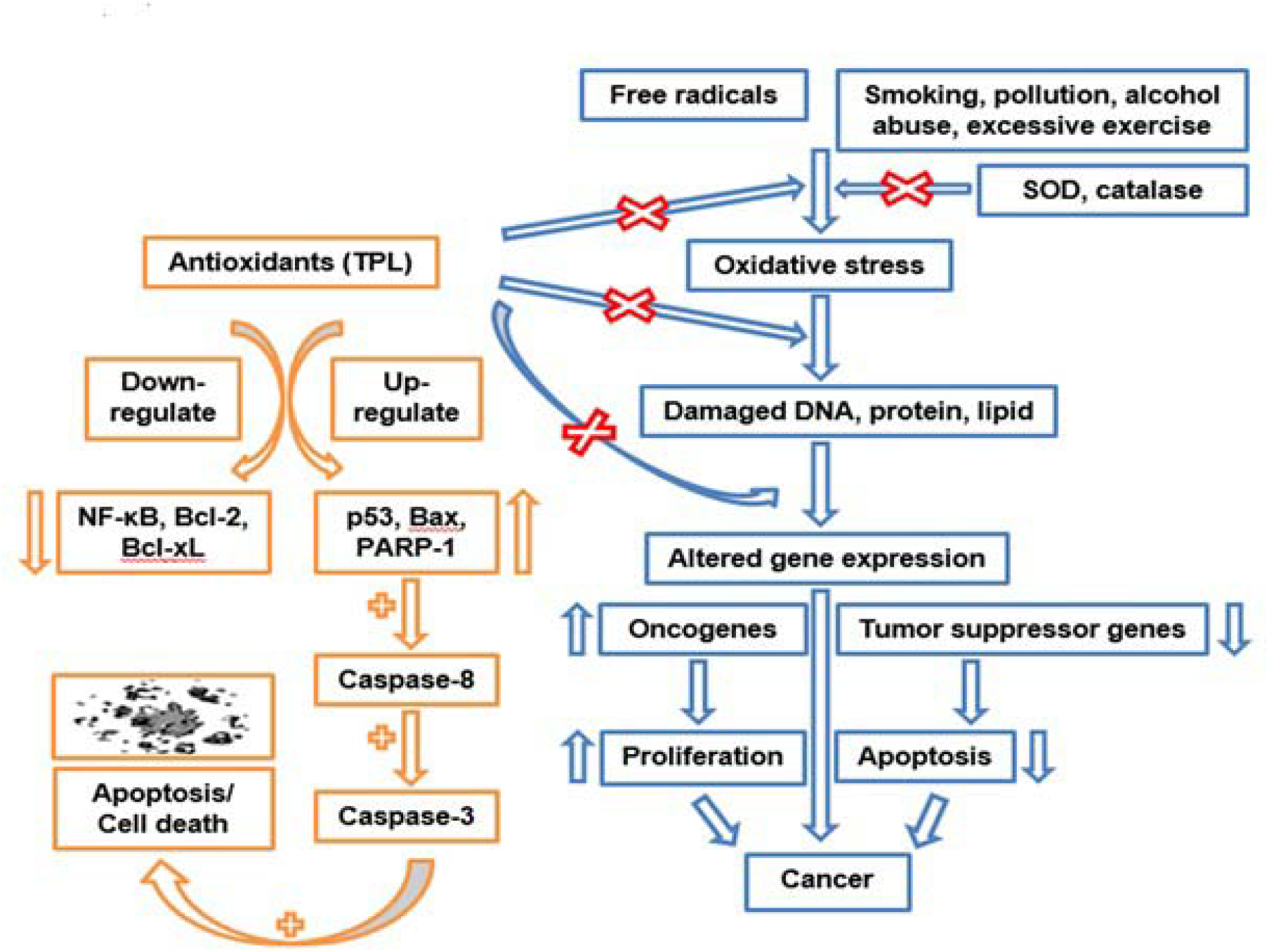

## 1. Introduction

Throughout the globe, causes of death due to cancer are increasing. This is often because of lack of early detection methods and poor prognosis with diagnosis at an advanced rather than an early stage [1]. According to the GLOBOCAN 2018 database, approximately 18.1 million new cancers were diagnosed and 9.6 million cancer deaths occurred in 2018. The WHO estimates that over 29.5 million new cases of cancer will be diagnosed annually, and deaths from cancer will be over 16.5 million a year by 2040 worldwide [2, 3]. Although cancer incidence rates are high in both developed and developing countries, mortality rates due to cancer are extremely high in developing countries. It was estimated that, in 2012, over 65% of all cancer deaths throughout the world occurred in lower and middle income countries and the percentage is projected to increase to 75% by 2030 [2, 4]. In the USA, approximately 1,762,450 cancer cases were diagnosed in 2019, which is the equivalent of more than 4,800 new cases every day. It was estimated that over 606,880 Americans died from cancer in 2019, representing around 1,700 deaths per day [5].This alarming rate of cancer mortality can be reduced with early cancer detection and effective anti-cancer medications.

The characteristics of cancer include abnormal cell proliferation and inhibition of, or resistance to, apoptosis [6, 7]. Apoptosis occurs through the regulation of distinct types of pro-apoptotic (e.g. Bax, Bid, Bak or Bad) and anti-apoptotic (e.g. Bcl-2, Bcl-xL, Mcl-1) genes [8]. In their review on cancer development, Hanahan and Weinberg reported that the apoptotic trigger occurs when there is an imbalance between pro- and anti-apoptotic genes, with the imbalance allowing uncontrolled proliferation of cells, cancer growth and progression [7]. The cellular characteristics of apoptosis include cell shrinkage and plasma membrane blebbing, chromatin condensation and DNA fragmentation [9]. The molecular characteristics include activation of caspases, changes in pro- and anti-apoptotic genes, and regulation of p53 and nuclear factor-κB (NFκB) signaling pathways [10-11]. Apoptosis occurs without any local inflammation, and is therefore the preferred mode of cell death in cancer elimination [12]. Therefore, if apoptosis is clearly understood, it opens a gate to tumor-specific apoptosis therapy.

Many natural products contain a variety of bio-active compounds. Their derivatives have been tested for anticancer potential on various preclinical cancer models [13]. From such a screening process, many anticancer drugs have been derived from plants, including vinblastine and vincristine from *Catharanthus roseus*, taxol from *Taxas brevifolia,* etoposide and teniposide from *Podophyllum species*, topotecan and irinotecan from *Camptotheca acuminate*, 4-ipomeanol from *Ipomoea bataatas*, and ^ß^-lapachone from *Tabebuia avellanedae*. These plant-based drugs induce apoptosis in various types of cancer [14, 15]. A major drawback of these drugs in cancer treatment is the indiscriminate killing of both the cancer and normal cells. Therefore, identification of effective anticancer drugs of plant origin with low toxicity to normal cells has gained renewed interest [16].

*Tabebuia pallida* (Lindl.) Miers (*T. pallida*), commonly known as white trumpet tree, belongs to Bignoniaceae family, and is a species of the genus *Tabebuia. T. pallida* is widely distributed in northern Mexico, Southern to Northern Argentina, the Caribbean Islands, Cuba, Chile and Paraguay [17]. Although our recent studies on *T. pallida* showed antimicrobial, antioxidant and anticancer activities [18-20], much remains unknown about its mechanistic role in these activities. Two reviews of multiple *Tabebuia* species demonstrated the anticancer efficacy against different types of cancers [21, 22]. For instance, the heartwood of *T. avellanedae* [15], leaves of *T. impetiginosa* [23], and leaves and flowers of *T. rosea* [24], have been used successfully against liver, breast, prostate and ovarian cancers. The stem bark extract of *T. avellanedae* and *T. cassinoides* had anticancer activity against Ehrlich Ascites Carcinoma (EAC) bearing mice (EAC mice) [25]. Moreover, β-lapachone, isolated from different *Tabebuia* species, was reported to act as an anticancer agent in a wide variety of tumor cell lines [22, 26].

The reported *in vitro* and *in vivo* biological activities of different *Tabebuia* species encouraged us to test the mechanism of anticancer activity of different parts (leaves, stem, root bark and flowers) of *T. pallida* in EAC mice. Already our group reported that *T. pallida* leaves (TPL) possess the highest antioxidant activity [18] and show strong anticancer toxicity with significant suppression of tumor cell growth [20]. The current study aimed to ascertain whether or not these anticancer effects were due to the induction of apoptosis, by activating caspases (caspase-3 and 8) and regulating p53 and NFκ-B signaling, and to identify the bioactive properties present in the extract that are responsible for the anticancer effect.

## 2. Materials and Methods

### 2.1. Plant Collection

In May 2013, leaves, stem and root bark and flowers of *T. pallida* were collected from the University campus, Rajshahi University, Bangladesh. Prof. Dr. Mahabubur Rahman, Department of Botany of the same University identified the plant and a voucher specimen of all plant parts was stored at the Herbarium of the Botany Department (Voucher No. MN-03) and the National Herbarium of Bangladesh (Voucher No. 46736). Plant parts were washed in water and shade dried with periodic sun drying for several days. The dried materials were then powdered using a grinding machine (Tinytech, India) and preserved at room temperature until utilized.

### 2.2. Preparation of the Extract

1.5 L of 80 % methanol was used to soak each powdered plant sample (500 g) for 7 days with periodic shaking and mixing. Filtration was performed using cotton then Whatman No.1 filter paper. The volume of extract was then reduced with a rotary evaporator (Bibby Sterlin Ltd, UK) under reduced pressure at 50 °C to obtain 30, 45, 40 and 35 g of extracted *T. pallida* leaves (TPL), stem bark (TPSB), root bark (TPRB) and flowers (TPF), respectively.

### 2.3. Chemicals

Taq polymerase, RNaseOUT, dNTPs, TRIzole reagent and Super Script-III reverse transcriptase were from Life Technologies (Invitrogen Bio Services India Pvt. Ltd, Bangalore, India). All other chemicals were of high grade.

### 2.4. Cytotoxicity by MTT Assay

The MTT (3-[4,5dimethylthiazol-2-yl]-2,5diphenyltetrazolium bromide) colorimetric technique was used to assess the cytotoxicity of TPL, TPF, TPSB and TPRB extracts against EAC cells [9]. In this experiment, EAC cells were collected from 7 day old EAC mice after 1 day of EAC intraperitoneal (i.p.) cell inoculation in healthy mice, and 1×10^6^ cells placed per well in 200 μL RPMI-1640 media in a 96-well plate. Five different concentrations (8 μg, 15 μg, 30 μg, 60 μg, and 120 μg/mL; N=3) of the extracts were added to each well of EAC cells to assess the maximal efficiency. As a control, dimethyl sulfoxide (DMSO), as the sample solvent (0.1% w/v), was utilized. The cells were then incubated at 37 °C for 24 h in a CO_2_ incubator and the MTT assay performed using routine procedures [27]. Subsequently, absorbance was measured at 570 nm. The cytotoxic effects of the extracts were estimated in terms of % growth inhibition and expressed as IC_50_ which is the concentration of the tested sample that reduces the absorbance of treated cells by 50% with reference to the control (untreated) cells. The ratio of cell proliferation inhibition was calculated using the following equation:

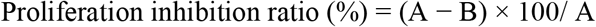

where A is the OD_570 nm_ of control solution and B is the OD_570 nm_ of sample solution.

### 2.5. In Vivo Antitumor Activity

#### 2.5.1. Experimental Animals

Four week old male mice *(Swiss albino)*, 20-25 g body weight, from the International Centre for Diarrheal Disease and Research in Bangladesh (ICDDRB) were housed in propylene cages using a 12 h dark and light cycle and temperature 25±2°C. Animals received food and water *ad libitum*. All mice were accustomed to the laboratory conditions for one week before experimentation. Animals were denied food and water for 12 h before the experiments.

#### 2.5.2. Ethics Approval

The Animal Ethical Committee, Rajshahi University (27/08/RUBCMB) and the Committee of Cell Research of Rajshahi Medical College, Bangladesh (ref. RMC/ER/2010-2013/01) approved the protocol of using mice as an animal model for cancer investigation.

#### 2.5.3. Cell Lines

Indian Institute of Chemical Biology (IICB), Kolkata, India provided the EAC cells that were cultured in *Swiss albino* mice as an ascites tumor by bi-weekly i.p. inoculation of 1×10^6^ cells/mouse.

#### 2.5.4. Experimental Design

There were 7 groups and in each group, there were 12 mice randomly divided. Group I was a control group receiving vehicle only; Group II was a tumor group receiving an i.p. injection of 1×10^6^ exponentially growing EAC cells; Groups III, IV, V, VI were tumor-bearing treated groups receiving an i.p. injection of 1×10^6^ exponentially growing EAC cells with TPL, TPSB, TPRB, TPF extracts at a dose of 100 mg/kg/day, respectively; and Group VII was a bleomycin-treated (0.3 mg/kg/day) tumor-bearing standard group. Exponentially growing EAC cells had been propagated i.p. (bi-weekly) in donor *Swiss albino* mice and collected from ascites fluid 6–7 days after inoculation of tumor cells [9].

#### 2.5.6. EAC Cell Administration

EAC cells, collected as above, were diluted in normal saline (0.9%) to 1×10^6^ cells/mL as measured by haemocytometry. Cell viability was determined by Trypan blue (0.4%) exclusion assay with results showing 99 % cell viability. Each tested *Swiss albino* mouse was administrated i.p. with 1×10^6^ of EAC cells in 0.1 mL saline.

#### 2.5.7. Determination of Toxic Effect of the Extracts

A total of 25, four week old male Swiss albino mice, 20-25 g body weight, were divided into five groups (n=5), and were used to assess any acute toxicity of the extracts. The test was performed with increasing oral dose concentrations of TPL, TPF, TPSB and TPRB in distilled water (50, 100, 200, 500 and 1,000 mg/kg body weight) that were administered orally at 20 mL/kg to each test group. The normal control group received distilled water (20 mL/kg). Following treatment, mice were allowed to feed *ad libitum* and were observed for 48 h for any mortality or behavioral changes [28].

#### 2.5.8. Determination of Inhibition of Cell Growth

*In vivo* inhibition of tumor cell growth was performed by a modification of the procedure reported by Sur and Ganguly [29]. In this method, 5 days of treatment were continued 24 h after tumor inoculation. Injection at volume 0.1 mL/day/mouse was given in each case. After transplantation of tumor cells, four mice out of 12 were sacrificed on the 6^th^ day and tumor cells collected with repeated i.p washing with 0.9 % normal saline. The Trypan blue exclusion method was performed to count the total number of viable tumor cells in the peritoneal cavity using the Cedex cell counter (Roche). Compared to the control (Group II), viable tumor cells of each mouse of the treatment groups were measured. The following formula was used to measure cell growth inhibition:

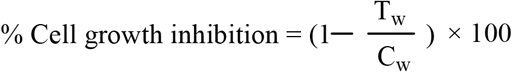

where T_w_ = mean number of tumor cells of Group III, IV, V, VI and VII and C_w_ = mean number of tumor cells of Group II.

#### 2.5.8. Determination of Average Survival Time and Tumor Weight

Similar conditions as mention in the previous experiment were adapted to measure these parameters. Weight change up to 20 days of treatment was monitored daily to assess tumor growth. According to the method described previously, host survival time was recorded and expressed as mean survival time in the days and percent increase of life span was calculated [30].

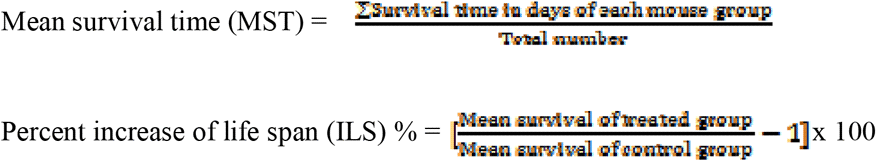

#### 2.5.9. Determination of Hematological Parameters

The hematological parameters (total RBC, WBC, and hemoglobin/Hb content) were measured by standard methods [31]. Blood was collected from the tail vein of each animal on the 12^th^ day after EAC cell inoculation. A hematometer was used to measure the Hb percentage and microscopy with a haemocytometer was used to estimate total WBC and RBC.

#### 2.5.10. Determination of Biochemical Parameters (Superoxide Dismutase and Catalase)

After the collection of blood samples on the 12^th^ day after EAC cell inoculation, the mice were sacrificed. The livers were excised, rinsed in ice-cold normal saline followed by cold 0.15 M Tris-HCl (pH 7.4), blotted dry, and weighed. A 10% w/v homogenate was prepared in 0.15 M Tris-HCL buffer followed by centrifugation at 1500 rpm for 15 min at 4°C. The supernatant thus obtained was used to estimate superoxide dismutase (SOD) and catalase (CAT) using standard procedures [32, 33].

#### 2.5.11. Examination of Morphological Change and Nuclear Damage

Cell apoptosis can be marked by plasma membrane blebbing, cell shrinkage, chromatin condensation and nuclear fragmentation (pyknosis), and chromosomal DNA fragmentation [9]. The apoptotic phenomena were observed morphologically by using a fluorescence microscope (OlympusiX71, Korea) as reported previously, using Hoechst 33342 staining [9]. In brief, EAC cells collected from Groups II, III, IV, V and VI were washed three times with phosphate buffer saline (PBS). The method of staining used 0.1 g/mL of Hoechst 33342 at 37 °C for 20 min in a dark room, and afterwards cells were washed once more with PBS.

#### 2.5.12. Measurement of Apoptosis

DAPI, Annexin V-fluorescein isothiocyanate (Annexin V-FITC) and propidium iodide (PI) triple fluorescent staining for cancer cell apoptosis was performed as described previously [27]. Briefly, 5×10^3^ cells per glass bottom dish were seeded and incubated overnight at 37 °C. After that, HeLa cells were treated with 100 µg/mL of TPL for 72 h at 37 °C. After aspiration, the cells were washed three times with 1X PBS followed by addition of 1 mL of 1X PBS per dish containing the staining solution of DAPI, Annexin V-FITC and PI in a ratio of 2:1:5. The dishes were incubated at 20-25 °C for 5 min under dark conditions, washed with 1X PBS twice and were observed under a confocal laser scanning microscope (Olympus, FV1000, Tokyo, Japan).

#### 2.5.13. Determination of Involvement of Caspases in T. pallida-Induced Apoptosis

To confirm the involvement of caspase-3 and −8 in the *T. pallida*-induced tumor cell death, untreated EAC cells in RPMI 1640 medium were incubated with z-DEVD-fmk (caspase-3 inhibitor, 2 µmol/mL) and z-IETD-fmk (caspase-8 inhibitor, 2 µmol/mL) for 2 h [34]. Then the cells were treated with 120 µg/mL of TPL and incubated at 37 °C in 5 % CO_2_ incubator for 24 h. Lastly, an MTT assay determined the cell growth inhibition.

#### 2.5.14. Isolation and Purification of Total RNA and Synthesis of cDNA

Total RNA was extracted using published methods [35]. In brief, EAC cells were collected from the peritoneal cavity of mice and washed with ice cold PBS twice followed by adding 0.5 mL TRIzole reagent (Invitrogen). TRIzole reagent treated EAC cells (1×10^6^ cells/mL) were gently vortexed for a few seconds. Then 0.2 mL CHCl_3_ was added with vigorous shaking and the mixture was incubated at RT for 2-3 min. This mixture was then centrifuged at 4 °C and 12,000 rpm for 15 min, the supernatant collected into an Eppendorf tube and 500 μL isopropanol added. The reaction mixture was then incubated for 10 min at room temperature (RT) and again centrifuged at 4 °C, 12,000 rpm, for 10 min. After removing the supernatant, the RNA was washed with 75% ethanol then centrifuged at 4 °C, 7,500 rpm, for 5 min. After removal of the supernatant, the RNA pellet was dried at RT, and mixed with DEPC-treated RNase free water. The concentration of RNA was estimated by nanodrop (Thermo scientific nanodrop 8000). For cDNA synthesis, 1 μg of purified RNA was dissolved with 0.5 μg of oligo (dT) primer (Promega), 0.5 mM dNTPs, 1× first-strand buffer, 5 mM DTT (dithiothreitol), 2 units of RNase OUT (40 unit/μL) and 10 units of Super Script III reverse transcriptase (200 unit/μL) (Invitrogen), to make a total volume of 20 μL. This reaction mixture was incubated at 50°C for 1 h, and the reaction terminated by heating at 70 °C for 15 min.

#### 2.5.15. Determination of mRNA Levels of Pro-and Anti-Apoptotic Genes by RT-PCR

As described previously, the expression levels of p53, Bax, PARP, Bcl-2, Bcl-XL and NFκ-B were determined by RT-PCR [35]. For PCR, specific primers were used to amplify the first-strand cDNA (Table 1). Reagents containing 1X Taq polymerase buffer, 25 pmol each of forward and reverse primers, 2.5 mM of each dNTPs and 0.25 U of platinum Taq polymerase (Tiangen, China) were used to prepare 25 µL reaction mixtures. The PCR products of tested genes and housekeeping GAPDH gene were electrophoresed in 1.5% agarose gels, then the gels were stained with ethidium bromide, and finally visualized by UV transilluminator (Vilber Lourmat).

**Table 1:**
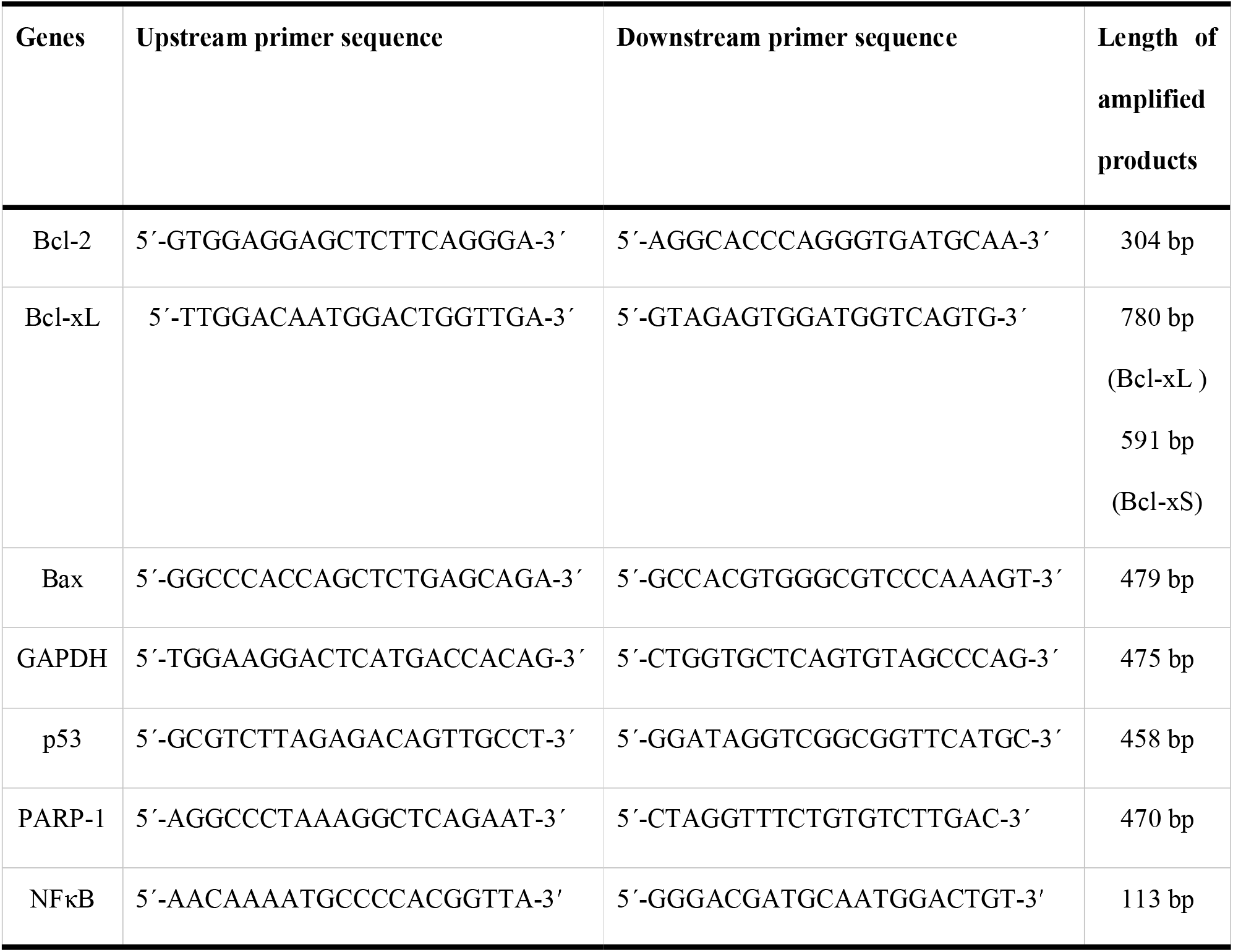
A list of primers with their upstream and downstream sequences and their subsequent length of amplified products

#### 2.5.16. Analysis by HPLC-PDA-ESI-MS/MS

The methanol extract of TPL (1 mg) was dissolved in a mixture of acetonitrile/water (50/50, v/v) before filtering through a PTFE 0.20µM syringe filter (Advantec, Tokyo, Japan). Then 2 µL filtrate was injected into the HPLC column. For conducting HPLC-PDA-ESI-MS analysis, a Thermo Scientific Surveyor HPLC and an LTQ linear ion trap mass spectrometer (MS; Thermo Scientific Inc. Waltham, MA, USA) with an electrospray ionization (ESI; Burker Esquire 3000 plus) interface at 40 °C, coupled with a photodiode-array detector (PDA) set at 280 nm. UV spectra were scanned from 200 to 700 nm. A Hypersil GOLD C18 column (100 x 2.1 mm; I.D., 1.9 µM) with eluent A (H_2_O plus 0.05% trifluoroacetic acid, TFA) and eluent B (acetonitrile plus 0.05% TFA) was used for chromatographic separation, where the flow rate was 200 µL/min followed by a gradient elution profile: 0-25 min, 90-40 % A; 25-35 min, 40-2 % A; 38-43 min, 2-40 % A; and 43-45 min, 40 % A. Both the positive and negative modes were considered for ESI experiments. The settings were as follows: mass range measured m/z 150-750; ion trap temperature, 275 oC; Em, 5.0 Kv; drying N_2_, 10 mL/min; nebulizing N_2_, 30 psi; sheath gas flow, 35U; capillary voltage, 12V; and collision gas; helium. Peak area investigation of the observed peaks was performed for primary quantitative analysis. Tentative structural identification was conducted by mass-selecting the ion of interest with a selected ion chromatogram of the individual species. MS^n^ spectra were obtained in auto MS^2^ mode (the product ion of the base peak was selected automatically as a precursor ion for the next stage MS) and provided additional structural information. Instrument operation and data collection were achieved with commercially available software Xcalibur (Finnigan Corp., San Jose, CA. USA).

#### 2.5.17. In silico Molecular Docking

##### 2.5.17.1. Protein Preparation

The 3D crystal structures of the active site of HeLa cell 5IAE receptor protein (PDB: 5IAE) [36] and urate oxidase (PDB: 1R4U) [37] were downloaded in PDB format from Protein Data Bank. Then, the structures were prepared and refined using the Protein Preparation Wizard of Maestro V11.1, N.Y, USA using OPLS3 Force Field [38, 39]. A detailed description has been incorporated in the supplementary documents.

##### 2.5.17.2. Ligand Preparation

Identified compounds (D-glucuronic acid, pelargonidin-3-O-glucoside, Quercetin-3-glucoside, Lapachol, Beta-lapachone) were collected from Pubchem databases. Ligprep 2.5 within the Schrödinger Suite 2015 utilizing OPLS_2005 force field was used to build the 3D model of the ligands. Various ionization states of the compounds were generated at pH 7.0 ± 2.0 using Epik 2.2 of Schrödinger Suite. Up to 32 possible stereoisomers were retained per ligand for analysis.

##### 2.5.17.3. Receptor Grid Generation

Receptor grids were calculated for the prepared proteins such that various ligand poses would bind within the predicted active site during docking. In Glide of Schrödinger Maestro V11.1, grids were generated keeping the default parameters of Van Der Waals scaling factor of 1.00 and charge cutoff of 0.25 subjected to OPLS3 Force Field. A detailed description has been incorporated in the supplementary documents.

##### 2.5.17.4. Glide Standard Precision (SP) Ligand Docking

SP flexible ligand docking was carried out in Glide [38] within which penalties were applied to non-cis/trans amide bonds. A Van der Waals scaling factor and partial charge cutoff of 0.80 and 0.15, respectively, were selected for the ligand atoms. Final scoring was performed on energy-minimized poses and displayed as the Glide score. A detailed description has been incorporated in the supplementary documents.

### 2.6. Statistical Analysis

Data are demonstrated as mean ± SD of N=3 for each experiment. The student’s unpaired t-test was performed to estimate significance between the test samples and control. To measure the significant differences among different groups, one-way analysis of variance (ANOVA) followed by a Dunnett’s post hoc test was used. *P* value <0.05 was considered as statistically significant. Free R-software version 2.15.1 (http://www.r-project.org/) and Microsoft Excel 2007 (Roselle, IL, USA) were used for the statistical and graphical evaluations.

## 3. Results

### 3.1. Cytotoxic Activity of T. pallida Extracts

The cytotoxic effect of methanol extracts of TPL, TPF, TPSB and TPRB against EAC cells was evaluated by MTT assay where incubation time was 24 h with 8-120 µg/mL concentrations. All the extracts possessed different dose-dependent cytotoxic activity against EAC cells. Viability of EAC cells was significantly inhibited by all extracts, with maximum inhibition by TPL (86.22%) compared to other extracts such as TPF (72.88%), TPSB (76.88%) and TPRB (62.66%) at the concentration of 120 µg/mL (Figure 1A). The IC_50_ (concentration producing 50% growth inhibition) of the TPL extract was 21.5 µg/mL, compared with IC_50_ values of 46, 37.5 and 57 µg/mL for TPF, TPSB and TPRB extracts, respectively (Figure 1B).

**Figure 1.**
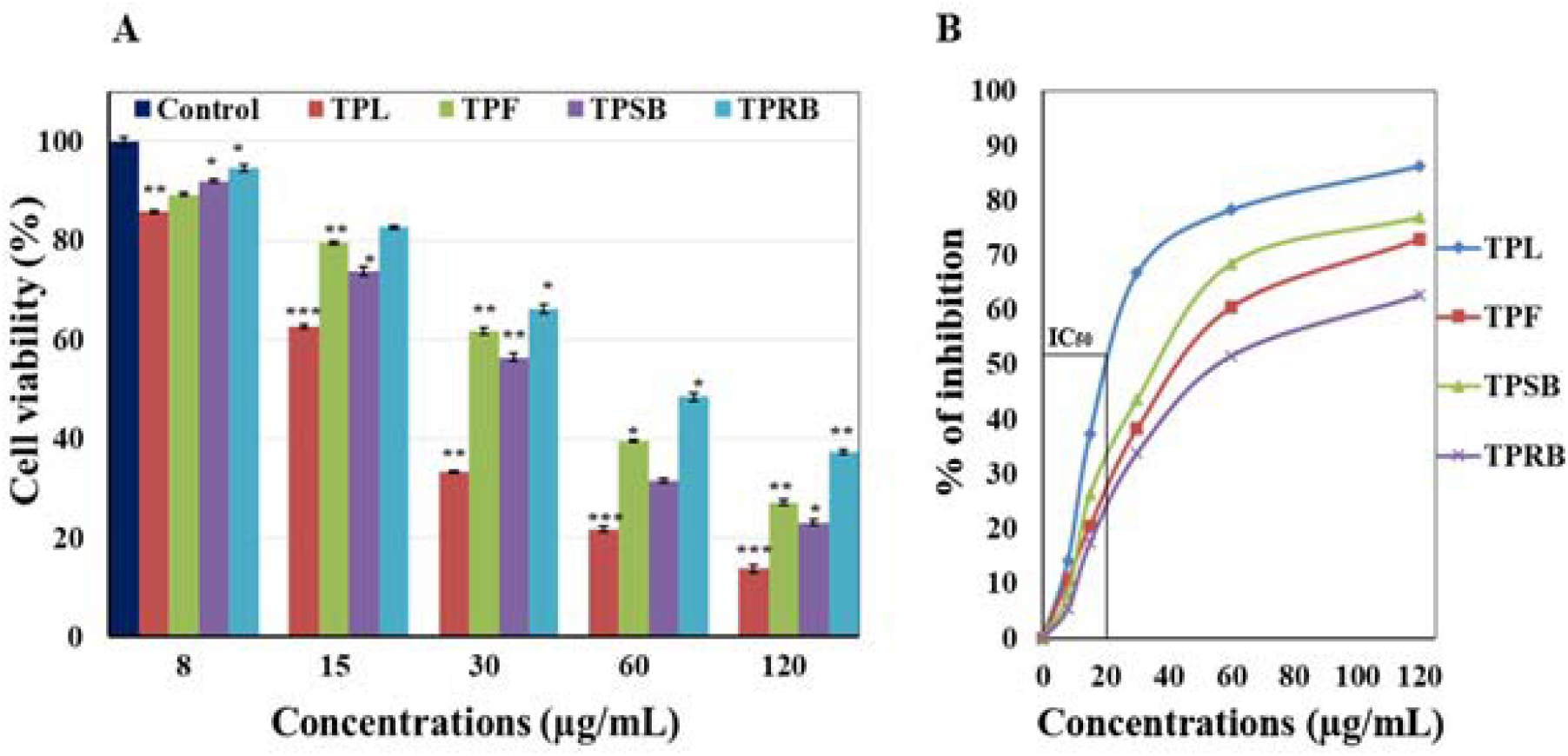
**(A)** Observation of EAC cells growth inhibition by *T. pallida* extracts. The cell proliferation was measured by the MTT assay (n = 3, mean ± SEM) after cells were treated with various doses of sample extracts and without samples for 24 h in CO_2_ incubator at 37 °C. ^∗^p<0.05,^∗∗^p<0.01 and ^∗∗∗^p<0.001 as compared sample-treated EAC cells with untreated control. **(B)** IC_50_ of *T. pallida* extracts (TPL, TPF, TPSB and TPRB) determined from EAC cell growth inhibition activity in MTT assay. Data expressed as mean ± SEM) (n = 3) for all tested dosages.

### 3.2. Acute Toxicity Studies of T. pallida Extracts on Normal Mice

The primary aim of the acute toxicity studies was to establish the therapeutic index (the ratio between the pharmacologically effective dose and the lethal dose) within the same strain and species [28]. *T. pallida* extracts were safe at doses as high as 1,000 mg/kg (p.o.) body weight. The behavior of the mice was observed for the first 3 h, then every 4 h during the next 48 h period. The extracts did not induce mortality, behavioral changes, locomotor ataxia and diarrhea or weight loss in mice during the 48 h observation period. Furthermore, food and water intake did not differ among the groups studied.

### 3.3. Effect of T. pallida Extracts on Survival Time and Average Tumor Weight

Tumor weight of EAC mice was calculated *in vivo* where treatment with TPL, TPF, TPSB and TPRB extracts at the dose of 100 mg/kg was continued for 20 consecutive days. Among the extracts, TPL treatment showed the greatest tumor weight reduction (6.16 ± 0.09 g), which was significant (*p* < 0.001) when compared with untreated EAC mice (22.4 ± 0.35 g). The reduction of tumor weight was also observed at various time points (Figure 2). Moreover, the tumor reduction capability of TPL closely resembled that of the standard bleomycin (6.89 ± 0.11 g) (Figure 3A). Since the extracts reduced the tumor weight, the effect of these extracts was then tested on the survival time in EAC mice. The mean survival time (MST) and life span of the untreated EAC mice (MST 18.03 ± 1.73 days) was increased with all extracts (Figure 3B).

**Figure 2.**
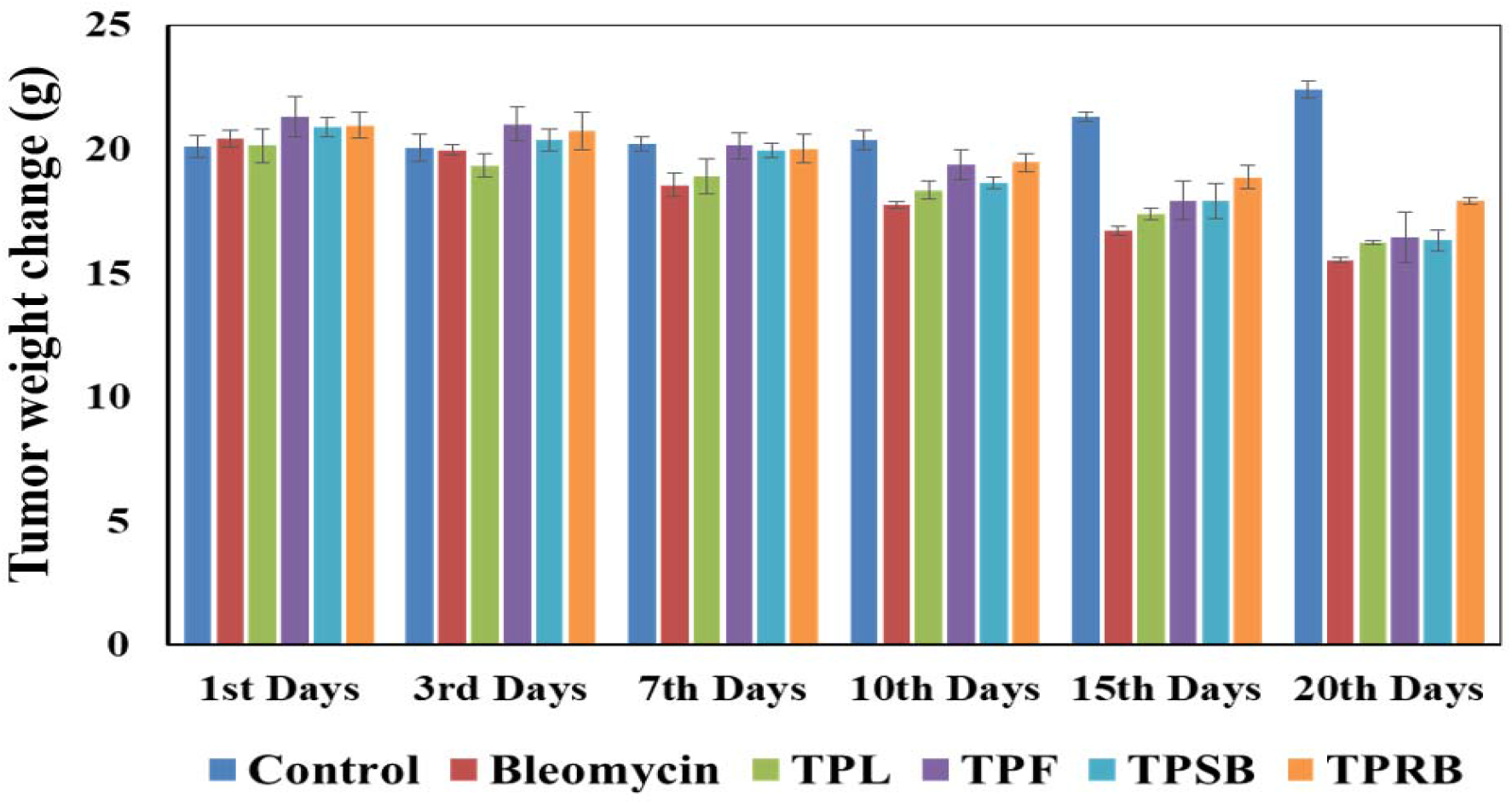
Effect of methanol extract of different parts of *T. pallida* on tumor weight of EAC cell bearing mice at different time points of the experiment. Values are mean ± SEM (n=4).

**Figure 3.**
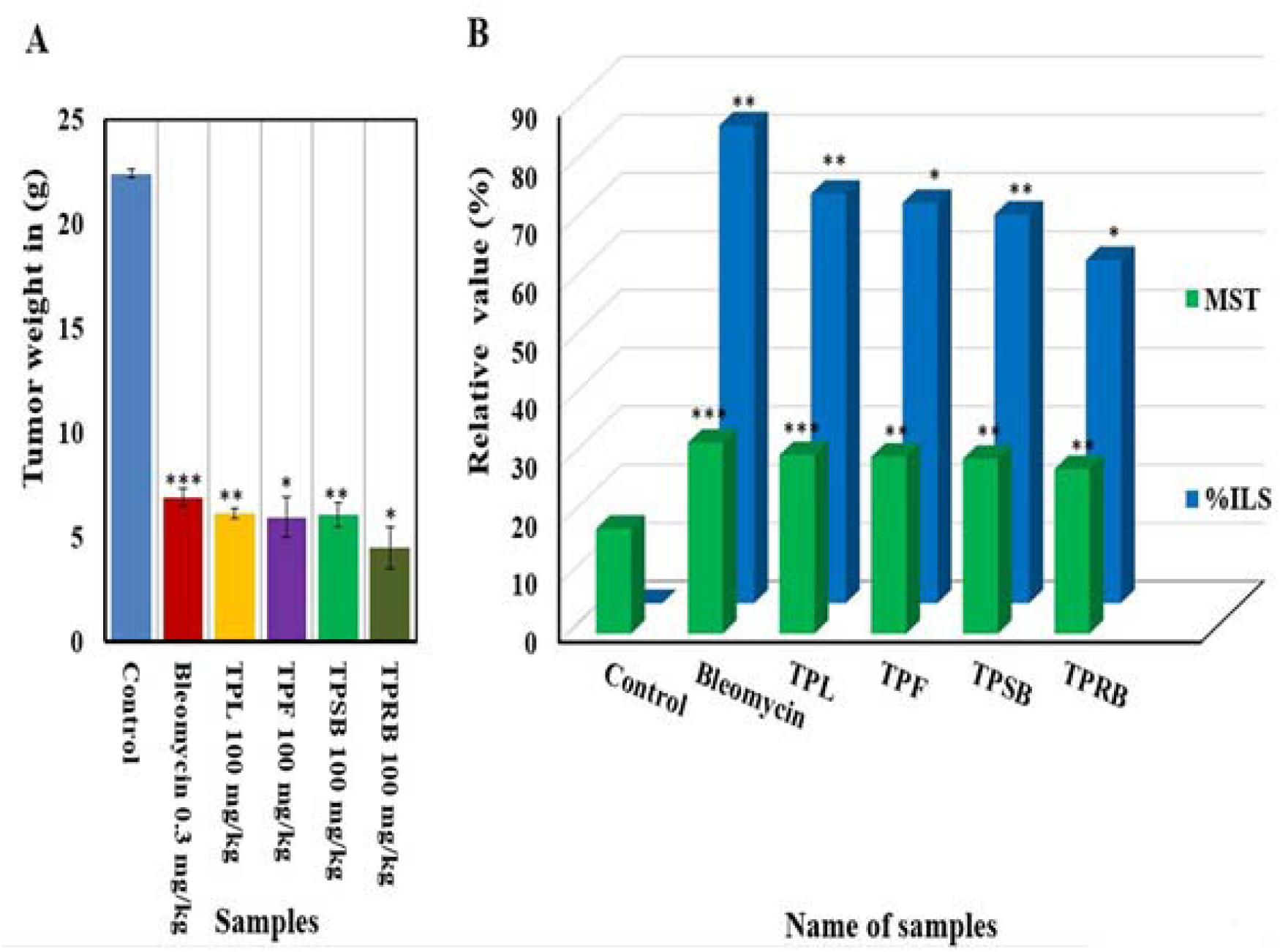
**(A)** Effect of methanol extract of different parts of *T. pallida* on tumor weight of EAC cell bearing mice. Values are mean ± SEM, (n=4), where significant values are, ^∗^p<0.05, ^∗∗^p<0.01 and ^∗∗∗^p<0.001 as compared to control. **(B)** Effect of methanol extract of *T. pallida* on mean survival time of EAC cell bearing mice and % increase of life span. Values are mean ± SEM, (n=4), where significant values are, ^∗^p<0.05, ^∗∗^p<0.01 and ^∗∗∗^p<0.001 as compared to control.

### 3.4. Effect of T. pallida Extracts on Hematological Parameters

The effect of the extracts on hematological parameters is presented in Table 2. The WBC and RBC count, and the percentage of Hb were significantly disrupted in EAC mice compared to the normal group. After 12 days of treatment with all extracts, there was a significant (*p*<0.05) improvement in the hematological parameters of EAC bearing mice with these parameters reverting to normal levels.

**Table 2:**
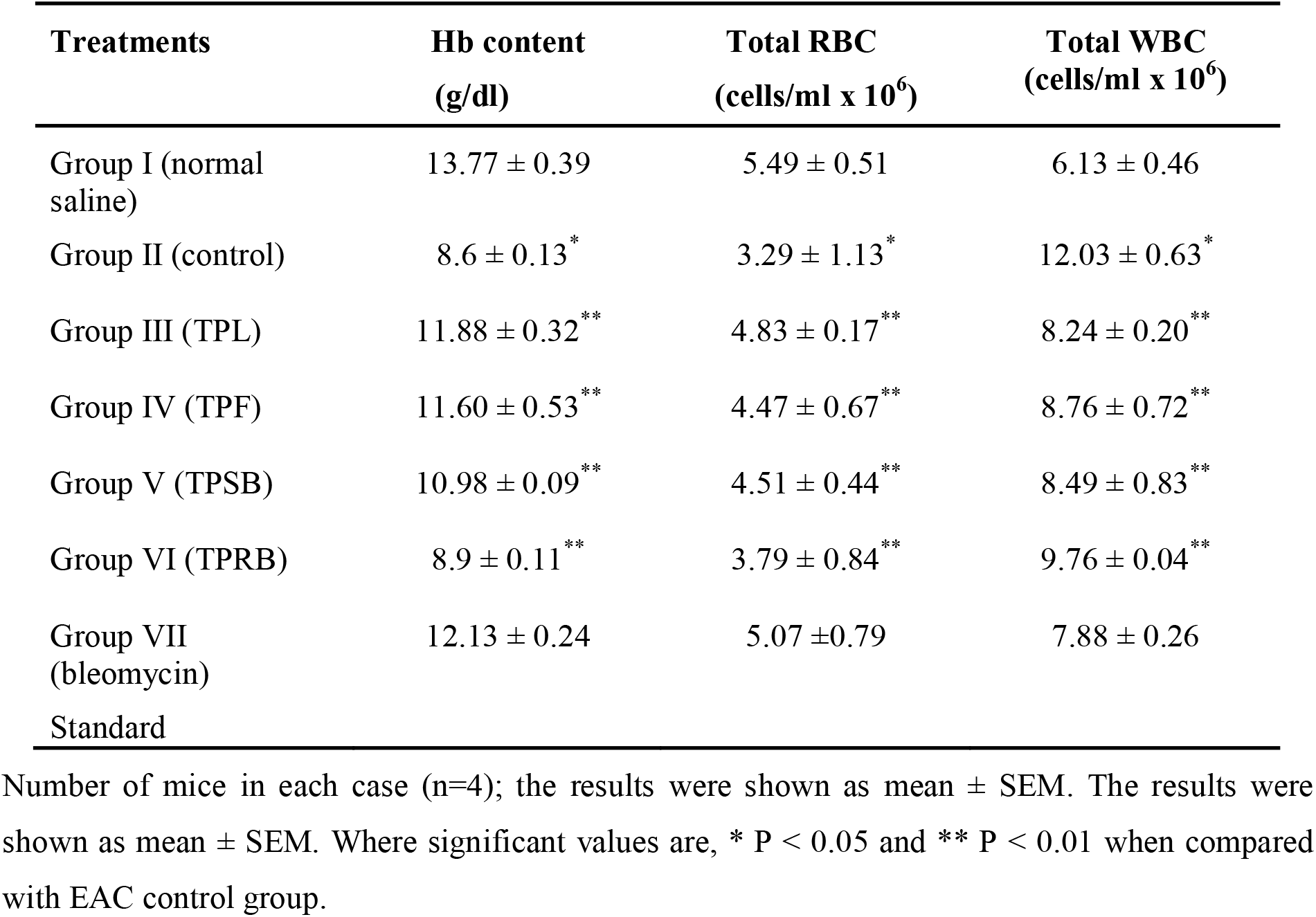
Effect of different parts of *T. pallida* extracts on hematological parameters in EAC mice.

### 3.5. Effect of T. pallida Extracts on Tumor Cell Growth

Plant derived extracts having phytochemical components are cytotoxic against tumor cells and show antitumor activity *in vivo* [8]. Since extracts from different parts of *T. pallida* confirm the presence of bioactive constituents with considerable *in vitro* cytotoxic effect, we investigated the *in vivo* effect of TPL, TPF, TPSB and TPRB on EAC mice. The TPL extract showed the greatest and significant (*p*<0.001) inhibition of tumor cell growth (71.71 %) in comparison to TPF (66.42 %), TPSB (63.13 %) and TPRB (57.62 %) at the dose of 100 mg/kg (i.p.). The standard bleomycin showed 83.10 % tumor cell growth inhibition at the dose of 0.3 mg/kg (i.p.) (Table 3). This result implies that extracts from the different *T. pallida* plant components, especially the TPL extract, had significant (*p*<0.001) anticancer activity compared to the standard bleomycin.

**Table 3:** Effect of different parts of *T. pallida* on EAC cell-induced tumor cell growth inhibition in mice.

| Name of Experiment | Nature of the drug | Dose mg/kg/day (i.p ) | No. of EAC cells in mice on day 6 after tumour cell inoculation | % of cell growth inhibition |
| --- | --- | --- | --- | --- |
| EAC cell bearing mice | Control | --- | $(5.41 \pm 0.40) \times 10^7^{**}$ | |
| Bleomycin | Standard | 0.3mg/kg | $(0.91 \pm 0.08) \times 10^7^{***}$ | 83.10 |
| TPL | Experimental | 100mg/kg | $(1.53 \pm 0.10) \times 10^7^{***}$ | 71.71 |
| TPF | Experimental | 100mg/kg | $(1.81 \pm 0.14) \times 10^7^*$ | 66.42 |
| TPSB | Experimental | 100mg/kg | $(1.83 \pm 0.17) \times 10^7^{**}$ | 63.13 |
| TPRB | Experimental | 100mg/kg | $(2.29 \pm 0.15) \times 10^7^*$ | 57.62 |

Number of mice in each case (n=4); the results were shown as mean ± SEM. Where significant values are, \* $P < 0.05$ , \*\* $P < 0.01$ and \*\*\* $P < 0.001$ when (EAC +TPL /TPF/ TPSB/ TPRB/ Standard) treated mice compared with EAC bearing control mice (EAC bearing only).

### 3.6. Effect of TPL Extract on Caspase-3 and −8 in TPL-Induced Apoptosis

We justified the selection of the TPL extract for further study because it was most effective in inhibiting EAC cell growth, and when the induction of apoptosis was measured using Hoechst stain and fluorescence microscopy, the characteristic morphological features of apoptosis of treated EAC cells were regularly identified when compared to controls (Figure 4). Caspase inhibitors, z-DEVD-fmk (caspase-3 inhibitor) and z-IETD-fmk (caspase-8 inhibitor), were used to check the effect of TPL on these caspases. TPL treatment caused 86.22 % suppression of tumor cell growth, which was significantly (*p*<0.01) reduced to 59.00 % and 66.00 % in the presence of caspase-3 and −8 inhibitors, respectively (Figure 5) suggesting that induction of apoptosis by TPL was caspase-3 and −8 dependent.

**Figure 4.**
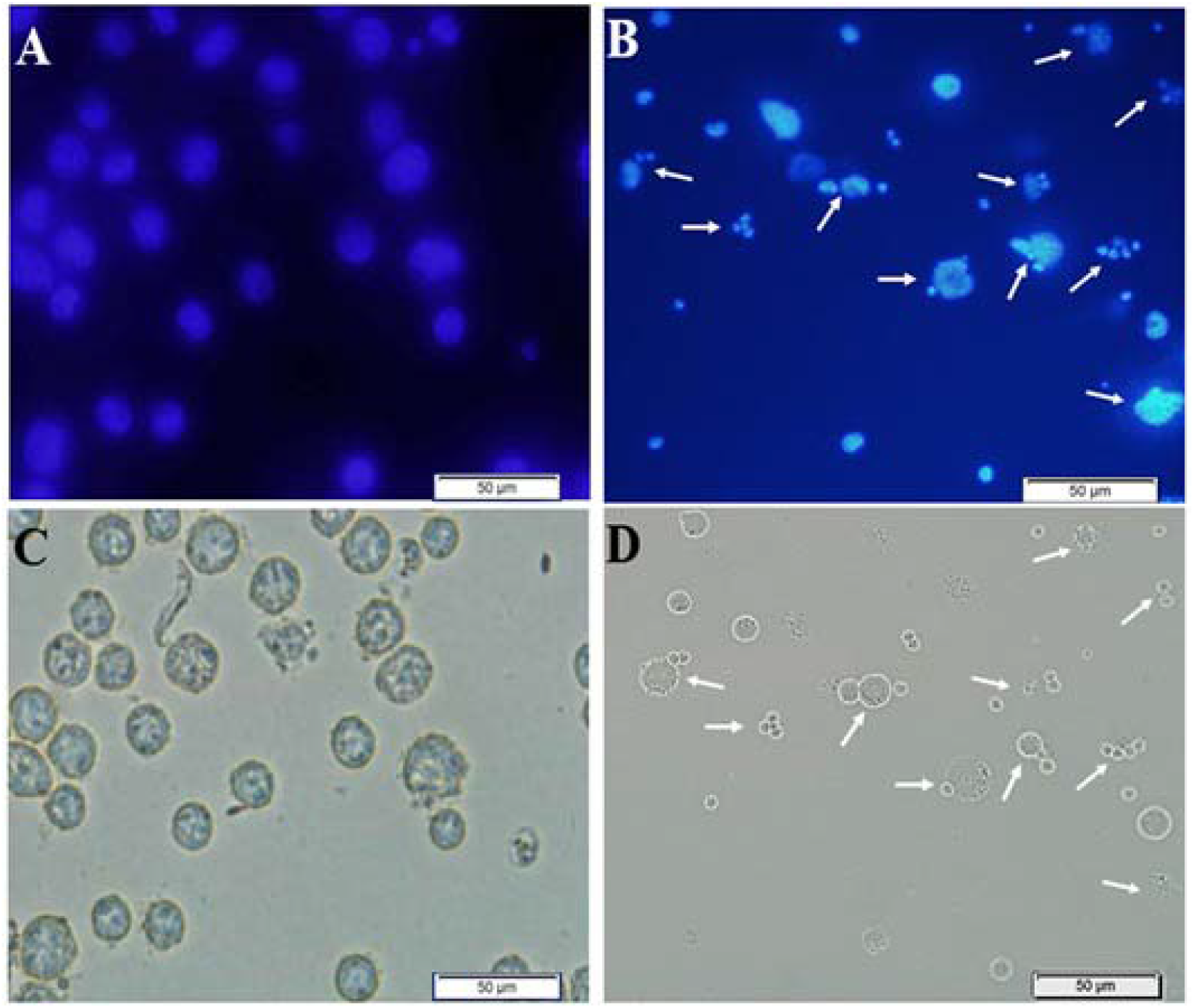
Morphological observation of TPL-induced apoptosis in Hoechst 33342 stained EAC cells by optical and fluorescence microscopy (Olympus iX71). Cells were collected from untreated EAC bearing mice demonstrated in A (fluorescence) and C (optical) and TPL treated EAC bearing mice demonstrated in B (fluorescence) and D (optical). Arrows indicate apoptotic features (condensed chromatin and nuclear fragmentation).

**Figure 5.**
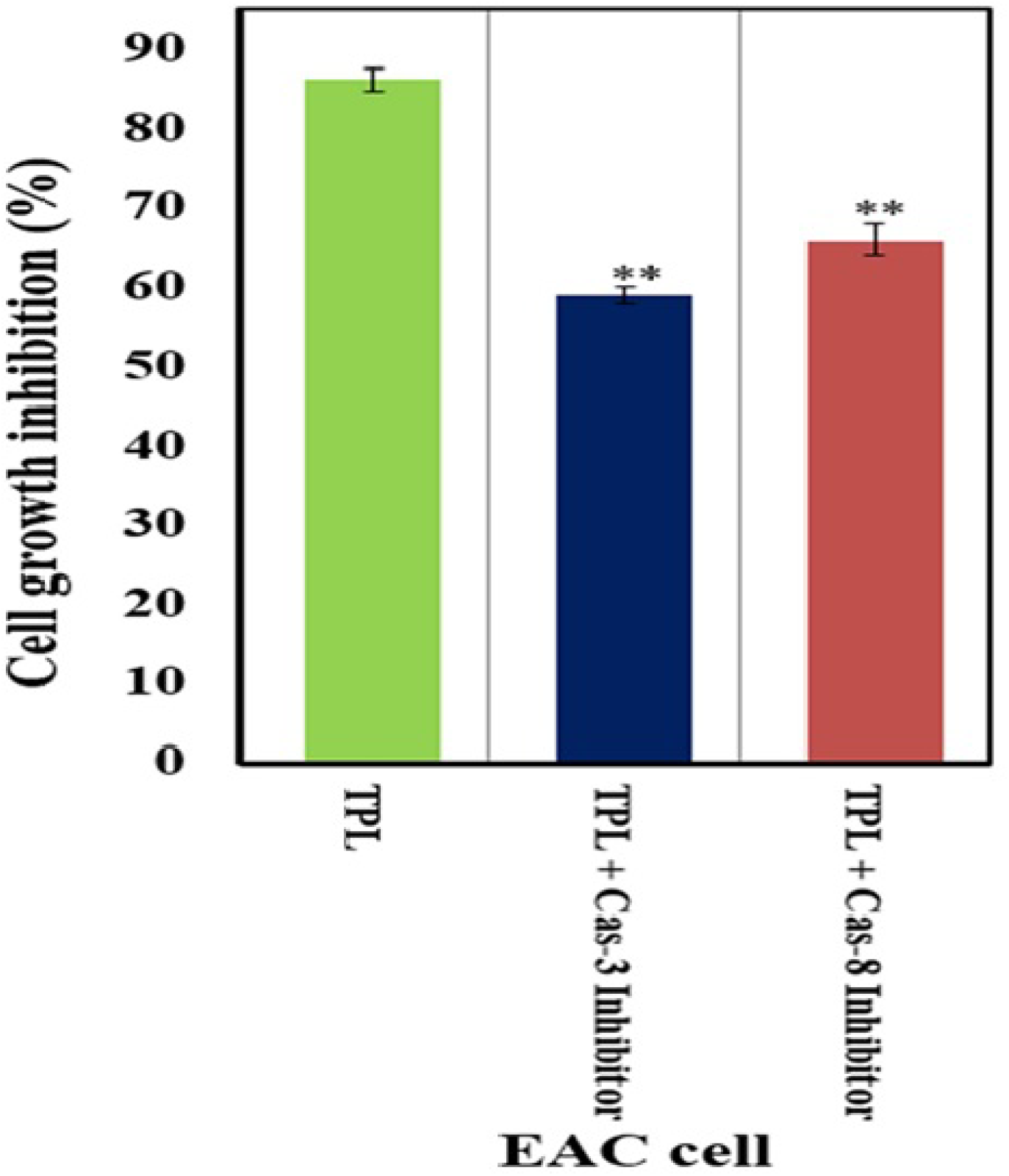
TPL induced apoptosis in a caspase-dependent manner. After pretreatment of the EAC cells with or without 2 mol/mL z-DEVD-fmk and z-IETD-fmk at 37 °C for 2 h, cells were incubated for 24 h at the said environment, then cell growth inhibition was determined by MTT assay (n = 3, mean ± SEM). Significant values are,^∗^P < 0.05, ^∗∗^P < 0.01 and ^∗∗∗^P < 0.001 when TPL treated EAC cells are compared with TPL + Cas-3/8 inhibitor treated EAC cells.

### 3.7. Effect of TPL on Cell Apoptosis

The apoptotic effect of TPL was further confirmed using HeLa cell apoptosis by evaluating DAPI, Annexin V-FITC, and PI triple fluorescence staining. DAPI is used as a marker of cell membrane permeability seen in very late apoptosis; whereas, Annexin V-FITC staining can identify apoptosis at an early stage [27]. In this study, DAPI, Annexin V-FITC, and PI signals could barely be detected in control cells (without treatment), while strong fluorescence densities were observed in response to treatment, indicating that TPL is able to induce HeLa cell apoptosis (Figure 6) through early and late stages.

**Figure 6.**
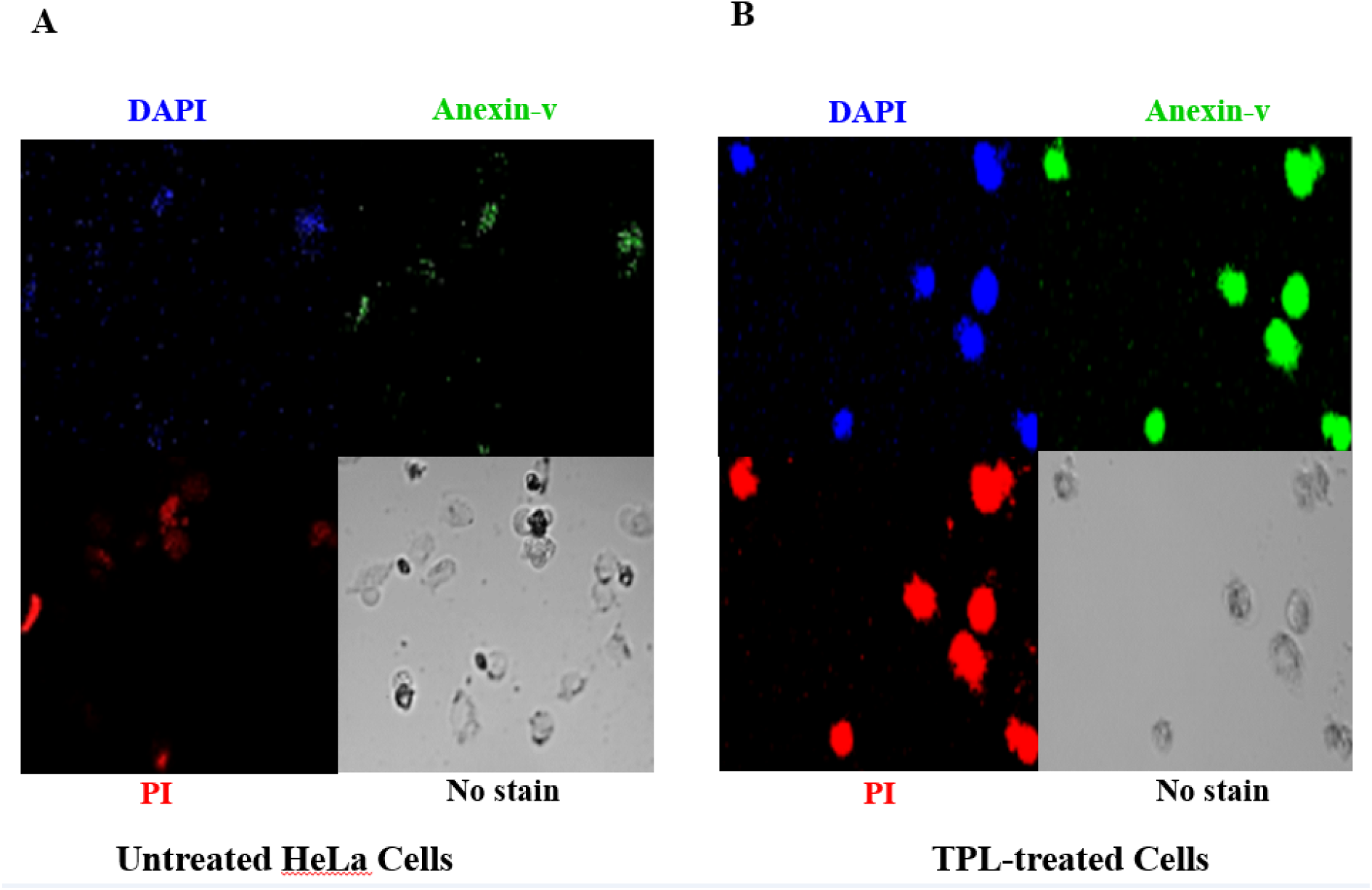
Representative images of DAPI, Annexin V-FITC and PI triple florescence staining on HeLa cell apoptosis. Cell nucleus was visualized by blue signal due to DAPI, Annexin V-FITCwas visualized by green signal, and PI was visualized by red signal.

### 3.9. Effect of TPL on Pro- and Anti-Apoptotic Gene Expression

Since it is widely reported that different pro-and anti-apoptotic genes play a vital role in induction of apoptosis, we therefore evaluated the involvement of TPL on anti-apoptotic genes (NFκ-B, Bcl-2, and Bcl-xL) and pro-apoptotic genes (p53, PARP-1, and Bax). In our study, TPL treatment significantly (*p*<0.05) increased the mRNA levels of pro-apoptotic p53, PARP-1, and Bax and decreased the expression of NFκB, Bcl-2 and Bcl-xL (Figure 7A, 7B, 7C). This observation confirmed that TPL treatment leads to a pro-apoptotic balance in pro- and anti-apoptotic genes, thereby promoting apoptosis of EAC cells.

**Figure 7.**
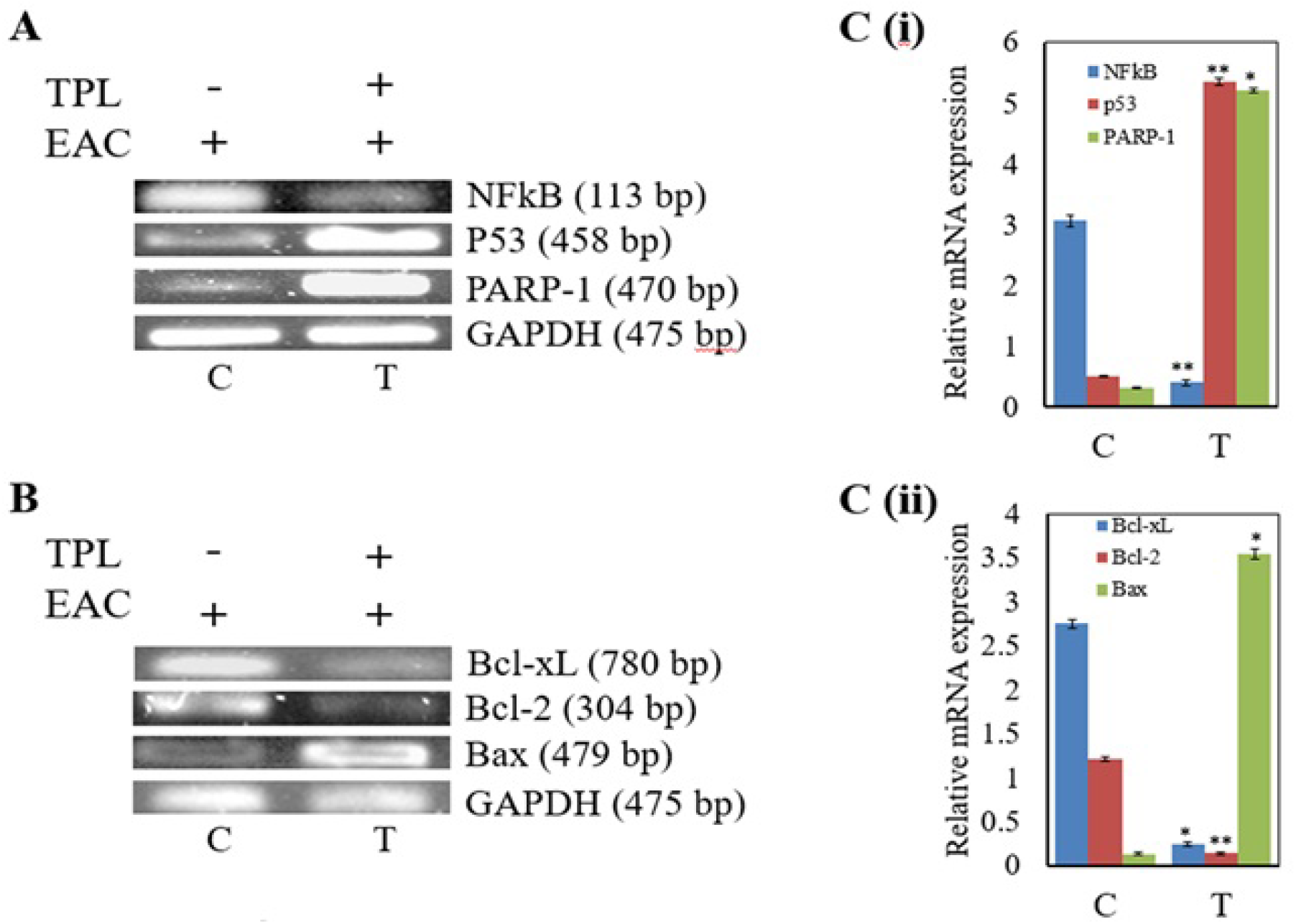
Analysis of mRNA expression levels of pro-and anti-apoptotic genes. Expression of (A) NFκ-B, p53, and PARP-1 (upper panel) and (B) Bcl-xL, Bcl-2, and Bax genes (lower panel) was analyzed by semi-quantitative RT-PCR in untreated EAC mice (control) and TPL-treated EAC mice. The positions of the genes along with their length are indicated on the left in bp. The bottom panel shows the PCR products of GAPDH as a control in both the panels. GAPDH transcript was used to normalize the expression levels. (C) (i), upper and (ii) lower panels indicate relative expression of (A, upper panel) NFkB, p53, and PARP-1 and (B, lower panel) Bcl-xL, Bcl-2, and Bax genes determined by a densitometric method. Error bars indicate the S.D. from three different experiments. C and T indicate untreated (control) and TPL-treated mice, respectively. The asterisks indicate that TPL treated tumor bearing mice is significantly different (^∗^p < 0.05, ^∗∗^p < 0.01, and ^∗∗∗^p < 0.001) from untreated control group.

### 3.10. Effect of TPL on Antioxidant Enzymes (SOD and CAT)

As shown in Figure 8A, SOD levels in the liver of EAC bearing mice significantly decreased by 37.41% in comparison with the non-EAC control mice (*p*<0.001). Treatment of EAC mice with TPL (100 mg/kg) increased SOD by 35.06%, as compared to that of the untreated EAC mice (*p*<0.01). The CAT levels in the EAC mice were significantly decreased by 59.1% in comparison with the non-EAC mice (*p*<0.001). Administration of TPL (100 mg/kg) to EAC mice increased the CAT levels by 50.22% when compared to the EAC control mice (*p*<0.01) (Figure 8B).

**Figure 8.**
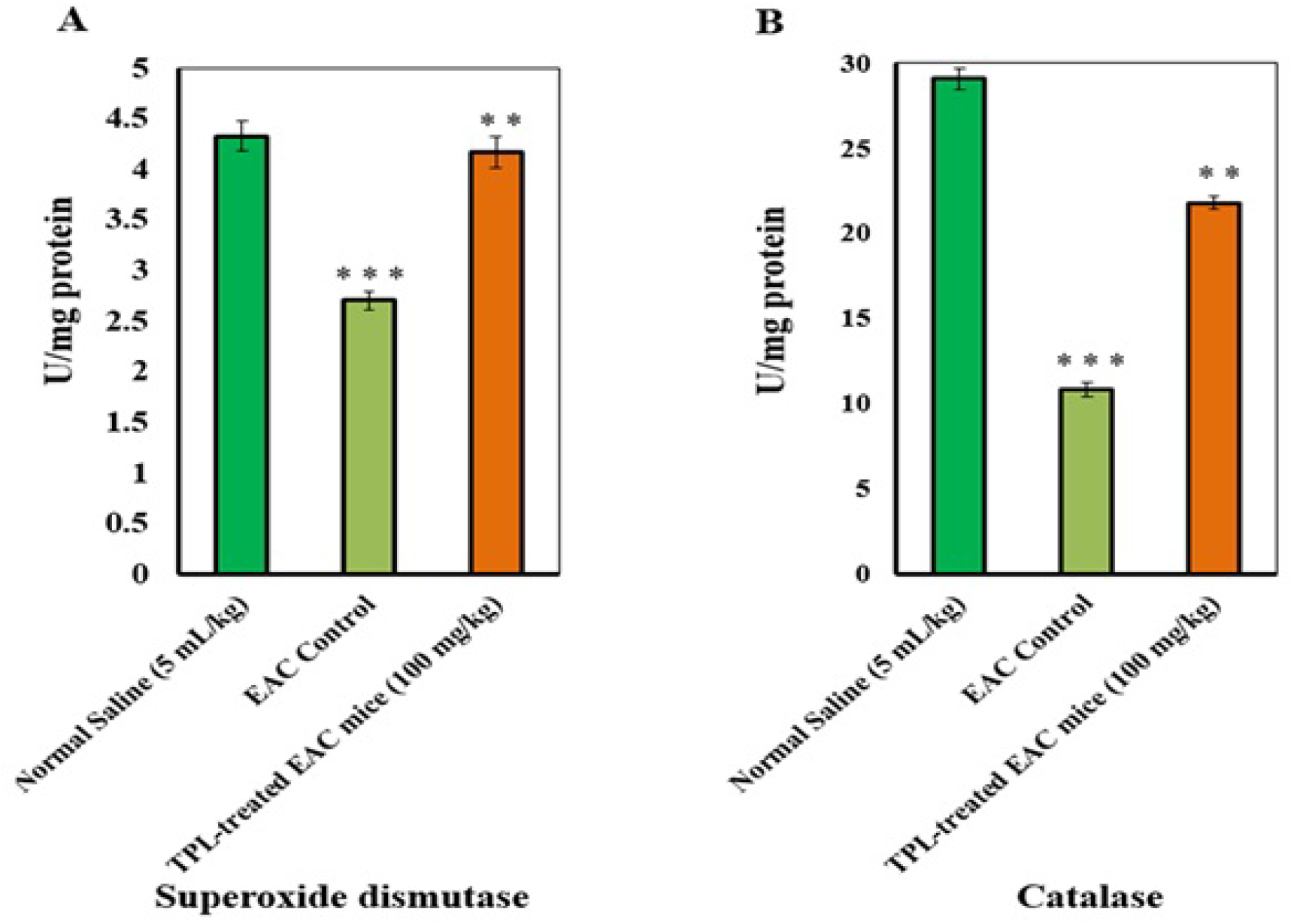
**(A).** Effect of the TPL on hepatic SOD activity in EAC mice. Data are expressed as the mean results in 4 mice ± SEM. ^∗∗∗^P<0.001, EAC control group compared to the normal group. ^∗∗^P<0.01, TPL-treated EAC mice compared to the EAC control group. **(B)** Effect of the TPL on hepatic catalase activity in EAC mice. Data are expressed as the mean results in 4 mice ± SEM. ^∗∗∗^P<0.001, EAC control group compared to the normal group. ^∗∗^P<0.01, TPL-treated EAC mice compared to the EAC control group.

### 3.11. Chemical Characterization of the TPL Extract by LC-PDA-ESI-MS/MS

A non-targeted LC-PAD-MS/MS chemical analysis of the highly active TPL extract was performed to identify the tentative components that might be considered to show the apoptosis leading anticancer effect. Figure 7A shows the major UV peaks of TPL at 280nm of HPLC spectrum. Compounds were tentatively identified based on parent molecular ions, UV values, retention times and elution order, as well as the fragmentation pattern described in the literature. The UV and MS data of the major compound, quercetin-3-glucoside (4), which were established on the basis of SIM analysis and by comparison with the published data [40], are shown in the Figure 9A and 9B. It was not possible to quantify the identified compounds in this mixture by LC/UV due to the overlapping and broad peaks, as well as a lack of corresponding commercial standards. Thus, the percentage of relative peak area of the tentatively identified compounds was used to determine the relative amounts of the constituents in TPL extract. The results indicated the presence of D-glucuronic acid, pelargonidin-3-O-glucoside, quercetin-3-glucoside, quercetin 3-O-(2’’-O-acetyl) glucoside, lapachol, β-lapachone, Coumarin glycoside and some unknown compounds (Table 4).

**Table 4:**
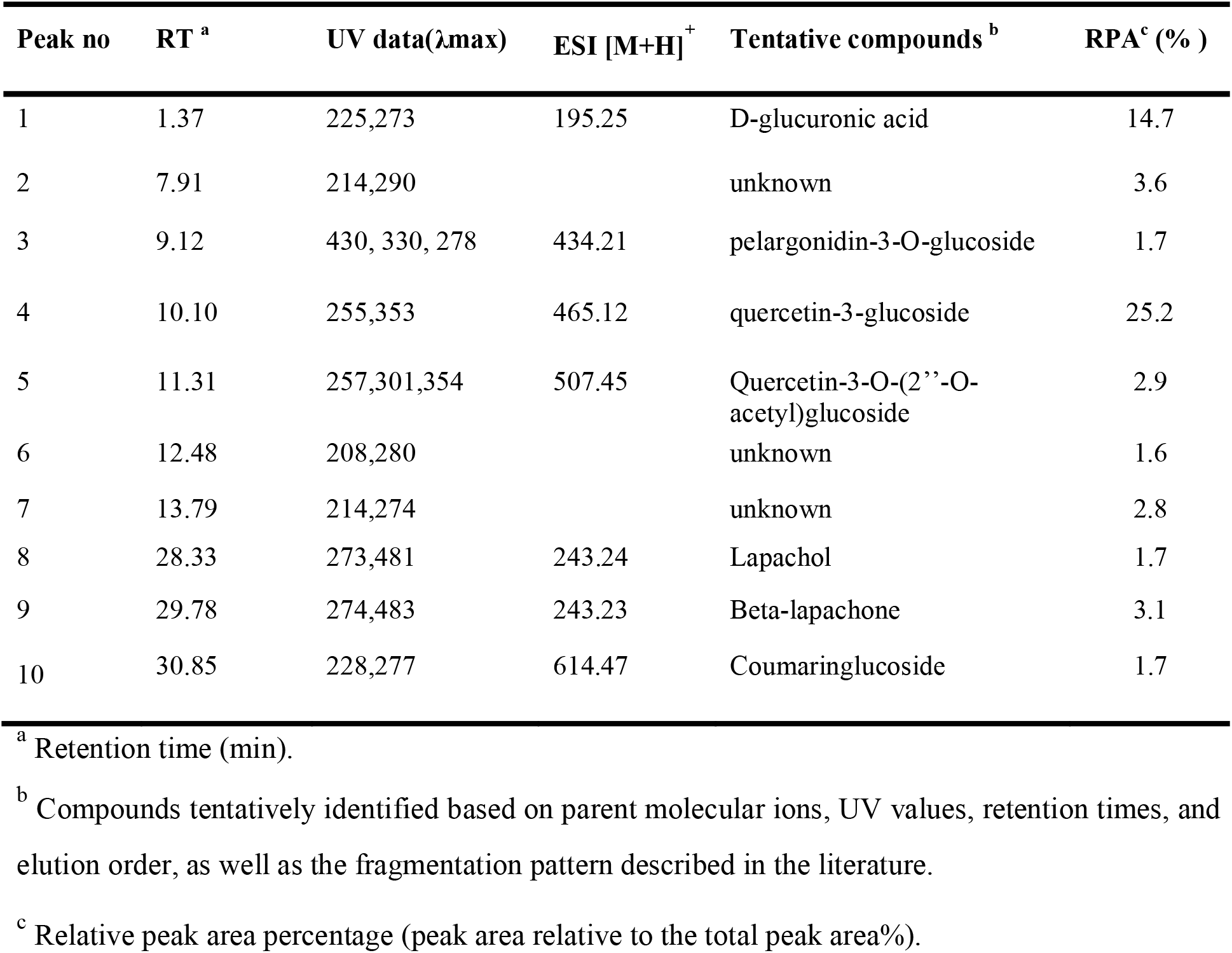
Chemical profiles of the major peaks and identified compounds in TPL.

| Peak no | RT <sup>a</sup> | UV data($\lambda$.max) | ESI [M+H] <sup>+</sup> | Tentative compounds <sup>b</sup> | RPA <sup>c</sup> ( % ) |
| --- | --- | --- | --- | --- | --- |
| 1 | 1.37 | 225,273 | 195.25 | D-glucuronic acid | 14.7 |
| 2 | 7.91 | 214,290 |  | unknown | 3.6 |
| 3 | 9.12 | 430, 330, 278 | 434.21 | pelargonidin-3-O-glucoside | 1.7 |
| 4 | 10.10 | 255,353 | 465.12 | quercetin-3-glucoside | 25.2 |
| 5 | 11.31 | 257,301,354 | 507.45 | Quercetin-3-O-(2''-O-acetyl)glucoside | 2.9 |
| 6 | 12.48 | 208,280 |  | unknown | 1.6 |
| 7 | 13.79 | 214,274 |  | unknown | 2.8 |
| 8 | 28.33 | 273,481 | 243.24 | Lapachol | 1.7 |
| 9 | 29.78 | 274,483 | 243.23 | Beta-lapachone | 3.1 |
| 10 | 30.85 | 228,277 | 614.47 | Coumaringlucoside | 1.7 |

<sup>a</sup> Retention time (min).
<sup>b</sup> Compounds tentatively identified based on parent molecular ions, UV values, retention times, and elution order, as well as the fragmentation pattern described in the literature.
<sup>c</sup> Relative peak area percentage (peak area relative to the total peak area%).

**Figure 9.**
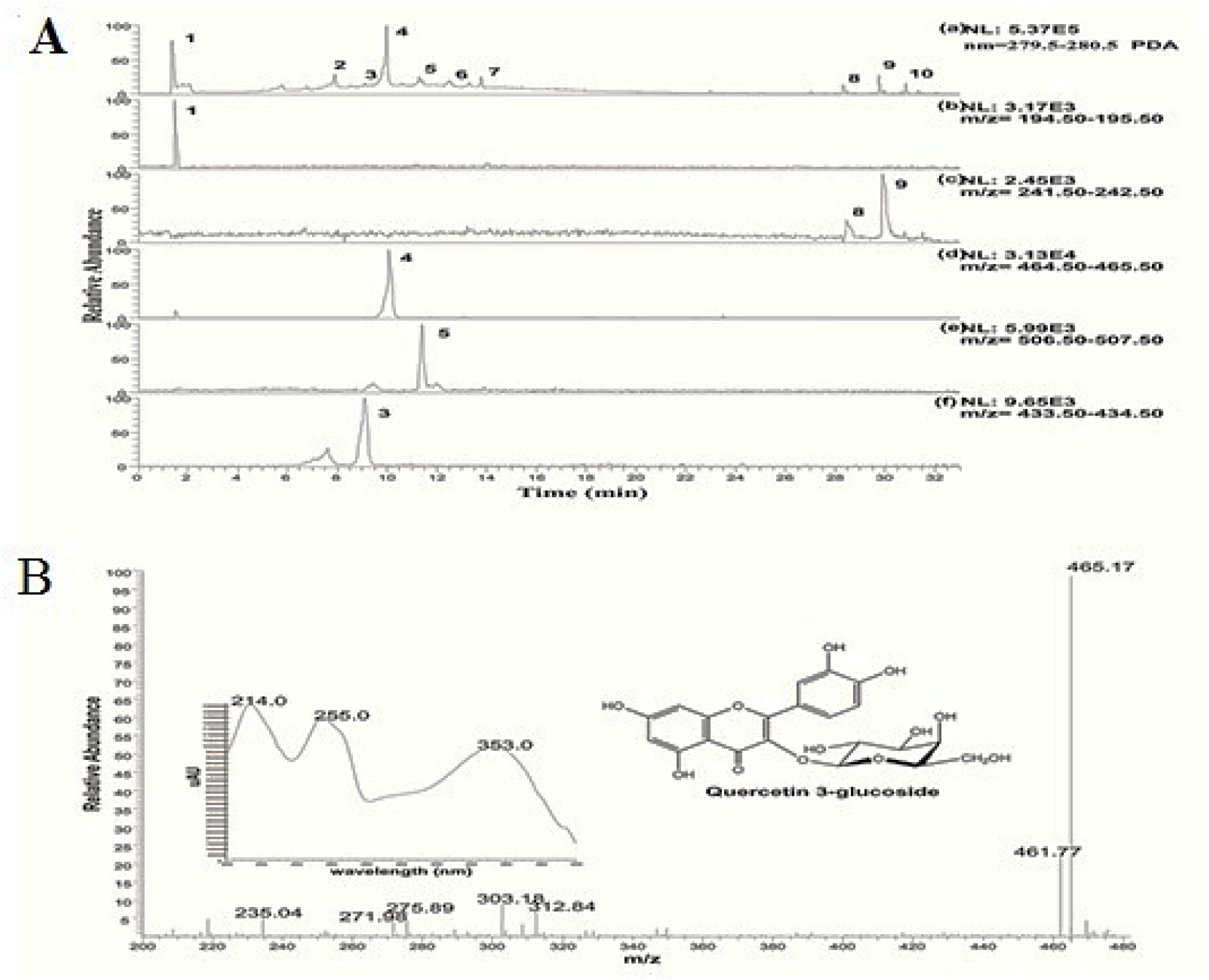
**(A)** HPLC profile of TPL. (a) HPLC-PDA spectrum of the major peaks at 280nm with tentatively identified compounds (1-10), (b-f) Selected ion chromatogram (parent ion) for the peak 8, 9, 4, 5 &3.**(B)** Mass spectrum of the major peak with UV spectra and structure of the quercetin 3-glucoside.

### 3.12. In silico Molecular Docking for Anticancer Activity of Compounds Present in TPL

Computer aided anticancer activity was performed to assess the binding pattern of molecules with the amino acids present in the active pocket of the protein. In this study, we used 3D crystal structures of the active site of the HeLa cell 5IAE receptor protein. Among the compounds obtained in TPL, pelargonidin-3-O-glucoside showed the highest docking score (−6.566) followed by D-glucuronic acid (−5.287), quercetin-3-glucoside (−5.238), beta-lapachone (−4.788), and lapachol (−4.092) with glide emodel (−59.813) and glide energy (−46.383) against active site of HeLa cell 5IAE receptor protein. The results of the docking analysis were depicted in Table 5 and Figure 10A-10E.

**Table 5.** Antioxidant and anticancer activities of identified compounds with binding protein PDB:1R4U and PDB: 5IAE.

| Compounds Name | 1R4U |  |  | 5IAE |  |  |
| --- | --- | --- | --- | --- | --- | --- |
|  | Docking score | Glide emodel | Glide energy | Docking score | Glide emodel | Glide energy |
| D-glucuronic acid | -5.54 | -35.243 | -25.4 | -5.287 | -43.462 | -30.721 |
| Pelargonidin-3-O-glucoside | -6.42 | -59.752 | -45.451 | -6.566 | -59.813 | -46.383 |
| Quercetin-3-glucoside | -5.396 | -53.77 | -42.743 | -5.238 | -57.228 | -45.294 |
| Lapachol | -4.678 | -38.195 | -30.876 | -4.092 | -32.423 | -26.351 |
| Beta-lapachone | -5.094 | -33.839 | -22.979 | -4.788 | -37.676 | -27.337 |

**Figure 10.**
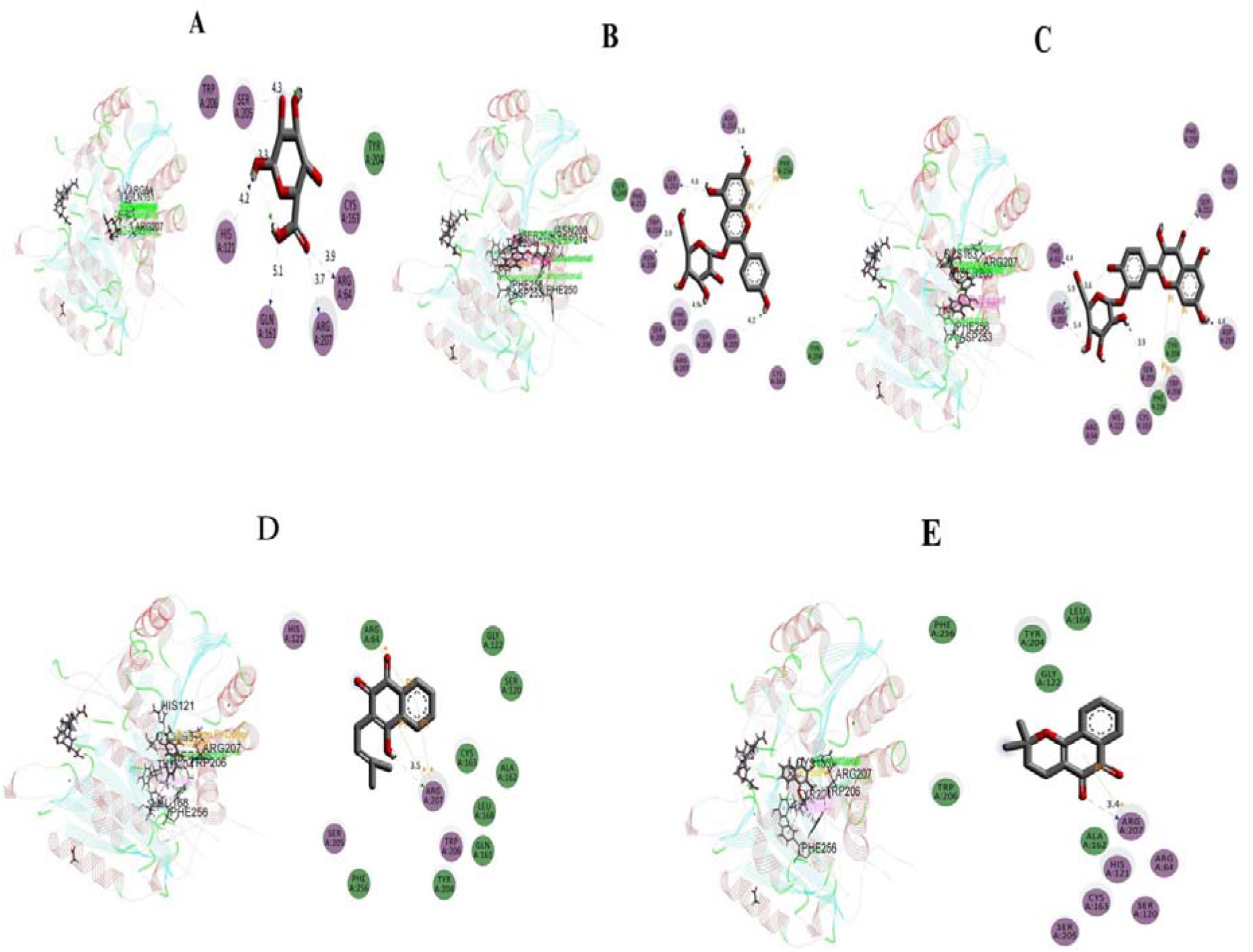
Best ranked poses and 2D interactions of **(A)** D-glucuronic acid **(B)** Pelargonidin-3-O- glucoside **(C)** Quercetin-3-glucoside **(D)** Lapacholand **(E)** Beta-lapachone with HeLa cell receptor (PDB: 5IAE) for anticancer activity.

The anticancer activity of the compounds was in the following order: pelargonidin-3-O-glucoside> D-glucuronic acid> quercetin-3-glucoside> beta-lapachone> lapachol.

### 3.13. In silico Molecular Docking for Antioxidant Activity

Computational study was performed to find out potential antioxidant molecules virtually. Grid and ligand based molecular docking program was used to assess the binding pattern of molecules with the amino acids present in the active pocket of the protein. The docking study was performed using the identified active compounds of TPL to the active site of urate oxidase (Uox) enzyme receptor (PDB: 1R4U). Pelargonidin-3-O-glucoside had the highest docking score (−6.42) with glide emodel (−59.752) and glide energy (−42.743) for binding with Uox (PDB: 1R4U) Table 5. The docking score of d-glucuronic acid, quercetin-3-glucoside, beta-lapachone, lapachol was −5.54, −5.396, − 5.094 and −4.678, respectively (Figure 11A-11E). The antioxidant activity of the compounds was in the following order: pelargonidin-3-O-glucoside> D-glucuronic acid> quercetin-3-glucoside> beta-lapachone> lapachol.

**Figure 11.**
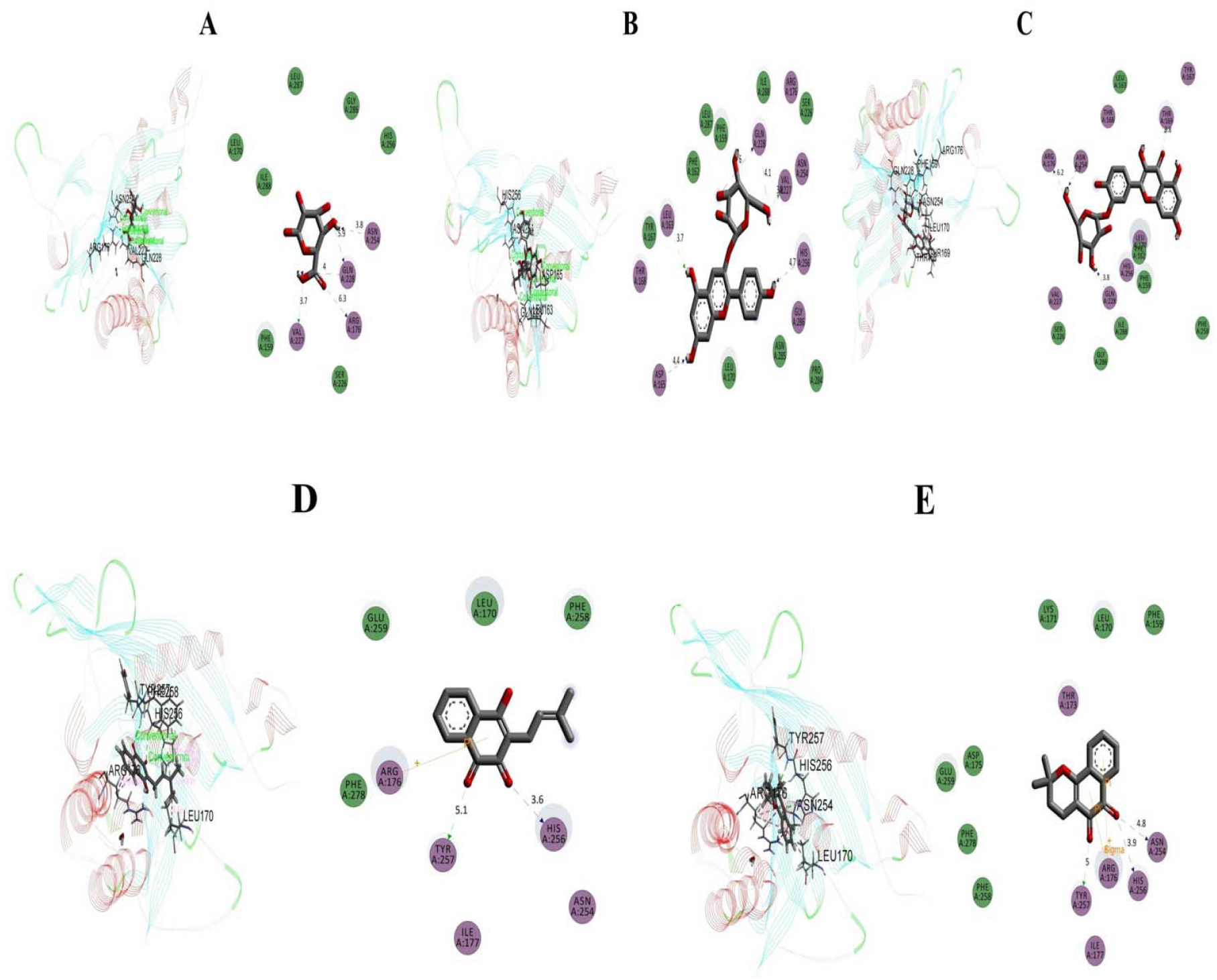
Best ranked poses and 2D interactions of **(A)** D-glucuronic acid **(B)** Pelargonidin-3-O- glucoside **(C)**Quercetin-3-glucoside**(D)** Lapacholand **(E)** Beta-lapachone with HeLa cell receptor (PDB: 5IAE) for anticancer activity.

## 4. Discussion

Cancer is a disease of unrestrained cell proliferation, where cancer cells develop the ability to escape apoptosis by utilizing various survival pathways [7]. Thus, in cancer therapy, the induction of apoptosis in cancer cells can be an effective treatment strategy. Inadequate effectiveness of existing chemotherapy drugs has lead to researchers investigating new apoptosis-inducing drugs to combat cancers.

Plant-derived natural products, crude extracts and/ or pure isolated compounds, have great potential in cancer treatment due to their capability to modulate apoptosis [41]. In our earlier observations, we confirmed that TPL contains the highest polyphenolic content, antioxidant and antiproliferative properties [18, 20]. The current study is an ongoing screening of our previous outcomes to determine whether the antioxidant enriched TPL possesses anticancer mechanisms at the molecular level in EAC mice. For the first time, this study provides evidence for the apoptosis-inducing capability of TPL and the LC-ESI-MS analysis of TPL extract for active content.

At first, various plant parts of *T. pallida* (TPL, TPF, TPSB and TPRB) were investigated for their cytotoxic activity. The TPL extract possessed significant cytotoxicity, presumably because of the presence of some highly active bioactive compounds. It has been reported that the phytochemical components existing in plants show antitumor activity via multiple anticancer pathways [42]. Moreover, cytotoxic compounds cause apoptosis by controlling signaling mechanisms such as activation of caspases under the regulation of the Bcl-2 family of proteins or upregulated expression of pro-apoptotic receptors on cancer cells, which causes activation of apoptotic signaling pathways by subsequent interaction with their ligands. Mitsuhashi et al. reported that phytochemicals such as polyphenolic compounds have cytotoxic and antioxidant properties [43]. Our previous findings of *in vitro* antioxidant activity of *T. pallida* support the current results of cytotoxic activity, especially of TPL extracts [18].

A reliable benchmark for judging the effectiveness of some antitumor drugs is the reduction of tumor cell growth, decreased tumor weight, decrease of WBC and the extension of life span of the treated animals [44]. Although all extracts of *T. pallida* had antitumor activity, the TPL extract was most active because it significantly suppressed the tumor cell growth, reduced the tumor weight and increased the life span of EAC mice.

Myelosuppression and anemia are major complications usually encountered in cancer chemotherapy due to decrease of RBC or the percentage of Hb in the host, resulting in iron deficiency in hemolytic or myelopathic conditions [45]. The treatment with different parts of *T. pallida*, especially TPL, improved Hb content and RBC levels, suggesting that these extracts possess protective action on the hemopoietic system.

Antioxidant enzymes such as SOD and CAT play an important role in maintaining the correct balance of free radicals in cells. These enzymes are involved in the clearance of superoxide and hydrogen peroxide (H_2_O_2_). SOD catalyses the attenuation of superoxide into H_2_O_2_, which is eliminated by CAT [46]. It has been reported that a decrease in SOD activity in EAC-bearing mice was due to loss of Mn^2+^ SOD activity in EAC cells and the loss of mitochondria, resulting in a decrease in total SOD activity in the liver. A small amount of CAT in tumor cells was reported [47]. Moreover, tumor growth is inhibited because of inhibition of SOD and CAT activities [32]. In the present study, similar findings were observed in EAC-bearing mice. The TPL treatment significantly increased the SOD and CAT levels. The *in vitro* [18], *in vivo* and *in silico* antioxidant activity of TPL was due to phytochemicals present in the TPL.

In addition, a compositional analysis of TPL (highly active part) was performed using LC-PDA-MS/MS to identify which phytochemicals are responsible for the anticancer activity. In the present study, the apoptotic activity of TPL could be attributed to the presence of quercetin-3-glucoside (major compound), pelargonidin-3-O-glucoside and/or lapachol derivatives, which were identified from TPL based on HPLC-MS evidence. The major phytochemical quercetin-3-glucoside and its derivatives and related compounds have well documented anticancer effects [48, 49]. Moreover, other compounds such as pelargonidin, lapachol and its derivatives also showed significant anticancer and pro-apoptotic effects against different cell lines [50, 51]. However, on the basis of the chemical profile, the methanol extract of TPL might also possess chemical entities or phytochemicals that interact among themselves, perhaps resulting in a synergistic effect, or in fact a cancelling effect, on their anticancer action. This needs further study. To corroborate the anticancer activity of TPL, *in silico* molecular docking was performed to obtain a good picture of antioxidant and anticancer compounds identified in *T. pallida*. In computer-aided drug design, *in silico* molecular docking is a pivotal tool which allows predicting the binding activity of compounds against particular proteins [52]. In addition, the possible molecular mechanism of actions of different pharmacological activities is determined comprehensively through molecular docking. In our study, a grid-based docking method was used to analyze the binding modes of the molecule with the amino acids present in the active pocket of the protein [53]. The interaction between the compounds and the active sites was assessed with docking analysis in Schrodinger Suite v 11.1. In this study, five major bioactive compounds of TPL interacted against two target receptors or enzymes, namely HeLa cell 5IAE receptor protein (PDB: 5IAE) and Uox enzyme receptor (PDB: 1R4U). Among these, pelargonidin-3-O-glucoside had the highest docking scores against Uox (− 6.42) and HeLa cell 5IAE receptor protein (−6.566). The negative and low value of the free binding energy (−6.42,-6.566) has proved to be a strong favorable connection with the respective receptors. From these results, the phytocompounds of *T. pallida* that were studied may, in part, be responsible for the anticancer and antioxidant activities through interactions with these target enzymes or receptors.

Cancer cells adapt different approaches to escape death, including modified expression of genes and proteins associated with cell life. To halt or stave off the apoptotic process is a common survival strategy of cancer cells, where deregulation of pro-apoptotic genes [54] or hyper-activation of anti-apoptotic genes takes place [7]. In our study, induction of apoptosis by TPL was confirmed by observing the alteration in nuclear morphology and cell shape in TPL-treated EAC cells compared to that of the untreated EAC cells. A series of caspases, such as caspase-3, −6, −7, −8 and −9, are involved in morphological changes and apoptotic cell death [55]. In this study, the activity of the caspase-3 and caspase-8 were blocked by the z-DEVD-fmk and z-IETD-fmk inhibitors, respectively. The results indicated that TPL induced caspase-dependent apoptosis by activating caspase-3 and caspase-8 significantly. In another apoptotic pathway, cellular commitment to apoptosis depends somewhat on the balance between proteins that intercede cell cycle arrest and cell death (e.g. p53, PARP-1, Bax) and proteins that are anti-apoptotic (e.g. Bcl-2 and Bcl-xL) [8, 56]. The pro-apoptotic gene, p53, is a key arbiter of apoptosis after DNA damage and cell cycle arrest [57]. Its activity subsequently allows Bcl-2 family members, particularly pro-apoptotic Bax, to pass an apoptotic signal to the mitochondria, eventually leading to cell death [58]. The current study indicated that there was p53 up-regulatio, down-regulation of Bcl-2 and Bcl-xL, and increased up-regulation of PARP-1 and Bax. As a result, the increased ratio of Bax/Bcl-2 consequently increased cell death. In a similar way, a decrease in NFκ-B signaling promoted apoptosis in association with decreased expression of Bcl-2 and Bcl-xL.

Furthermore, the effect of the TPL on cell apoptosis was evaluated using DAPI, Annexin V-FITC, and PI triple fluorescence staining [27]. In this study, DAPI, Annexin V-FITC, and PI signals were negligible in control cells (without treatment), while strong fluorescence densities were observed in response to treatment, verifying the capacity of TPL extractsto induce cell apoptosis.

Taking all the results together, we propose a mechanistic pathway of anticancer potential of TPL where antioxidant-enriched TPL blocks oxidative stress by scavenging free radicals [18]. This triggers apoptosis by regulating the inhibition of NFκ-B and its target genes Bcl-2 and Bcl-xL and increasing p53 and its target genes PARP-1 and Bax. An increase in p53 ultimately activates caspase-3 and-8, which induces apoptotic cell death. Phytochemicals with antioxidant potential were identified in TPL. Collectively, these outcomes support the established folkloric value and popularity of *T. pallida*. Therefore, *T. pallida* can be considered as a potential candidate for possible therapeutic intervention in cancer.

## 5. Conclusions

In this study using a pre-clinical mouse model, the TPL extract of *T. pallida* showed anticancer activity by inhibiting the growth of tumor cells. The TPL treatment-induced apoptosis was associated with activation of caspases-3 and −8, and was perhaps mediated by the regulation of p53 and NFκ-B. The LC-MS/MS profiling of TPL indicated the presence of different bioactive compounds, some of which were antioxidant, with a long history of use as anticancer drugs. In light of these data, TPL contains apoptotic pathway-inducing molecules. Computational study of the isolated compound from TPL revealed favorable anticancer and antioxidant activities through interactions with target enzymes or receptors. A more extensive study is needed to identify individual mechanisms of action of precise bioactives of the TPL extract for their anti-apoptotic application in cancer therapy.

## Acknowledgments

The authors are grateful to IICB, Kolkata, India, for providing the EAC cells, Central Science Laboratory of Rajshahi University, Bangladesh for giving support in fluorescence microscopy and also to ICDDRB, for supplying the experimental mice with standard mice pellets. We also thank Dr. AHM Mahabubur Rahman, Professor, Department of Botany, University of Rajshahi and National Herbarium, Dhaka, Bangladesh for the identification of the plant. The authors want to thank Dr. Syed Rashel Kabir, Professor, Department of Biochemistry & Molecular Biology, for his support in MTT assay.

